# The giant diploid faba genome unlocks variation in a global protein crop

**DOI:** 10.1101/2022.09.23.509015

**Authors:** Murukarthick Jayakodi, Agnieszka A. Golicz, Jonathan Kreplak, Lavinia I. Fechete, Deepti Angra, Petr Bednář, Elesandro Bornhofen, Hailin Zhang, Raphaël Boussageon, Sukhjiwan Kaur, Kwok Cheung, Jana Čížková, Heidrun Gundlach, Asis Hallab, Baptiste Imbert, Gabriel Keeble-Gagnère, Andrea Koblížková, Lucie Kobrlová, Petra Krejčí, Troels W. Mouritzen, Pavel Neumann, Marcin Nadzieja, Linda Kærgaard Nielsen, Petr Novák, Jihad Orabi, Sudharsan Padmarasu, Tom Robertson-Shersby-Harvie, Laura Ávila Robledillo, Andrea Schiemann, Jaakko Tanskanen, Petri Törönen, Ahmed O. Warsame, Alexander H.J. Wittenberg, Axel Himmelbach, Grégoire Aubert, Pierre-Emmanuel Courty, Jaroslav Doležel, Liisa U. Holm, Luc L. Janss, Hamid Khazaei, Jiří Macas, Martin Mascher, Petr Smýkal, Rod J. Snowdon, Nils Stein, Frederick L. Stoddard, Nadim Tayeh, Ana M. Torres, Björn Usadel, Ingo Schubert, Donal Martin O’Sullivan, Alan H. Schulman, Stig Uggerhøj Andersen

## Abstract

Increasing the proportion of locally produced plant protein in currently meat-rich diets could substantially reduce greenhouse gas emission and loss of biodiversity. However, plant protein production is hampered by the lack of a cool-season legume equivalent to soybean in agronomic value. Faba bean (*Vicia faba* L.) has a high yield potential and is well-suited for cultivation in temperate regions, but genomic resources are scarce. Here, we report a high-quality chromosome-scale assembly of the faba bean genome and show that it has grown to a massive 13 Gb in size through an imbalance between the rates of amplification and elimination of retrotransposons and satellite repeats. Genes and recombination events are evenly dispersed across chromosomes and the gene space is remarkably compact considering the genome size, though with significant copy number variation driven by tandem duplication. Demonstrating practical application of the genome sequence, we develop a targeted genotyping assay and use high-resolution genome-wide association (GWA) analysis to dissect the genetic basis of hilum colour. The resources presented constitute a genomics-based breeding platform for faba bean, enabling breeders and geneticists to accelerate improvement of sustainable protein production across Mediterranean, subtropical, and northern temperate agro-ecological zones.

## Main

Replacing meat or milk protein with plant-based alternatives, even partially, would make a large contribution to reducing carbon emissions^1^ and help to mitigate climate change. An obstacle to implementing this scheme is that meat-eating countries do not produce enough plant protein^2^. Common bean (*Phaseolus vulgaris*), chickpea (*Cicer arietinum*), and soybean (*Glycine max*), which supply dietary protein in tropical and subtropical countries, do not attain yield optima in temperate regions of Europe and America, where meat consumption is highest globally^3^. Faba bean (*Vicia faba* L., 2n=12) was domesticated in the Near East more than 10,000 BP^4, 5^ and its broad adaptability, value as a restorative crop in rotations and high nutritional density has propelled it to the status of a global crop grown on all continents except Antarctica^6^. It is the highest yielding of all grain legumes^7^ and has a favourable protein content (c. 29%) compared to other cool season pulses such as pea, lentil and chickpea, making it a suitable candidate to meet challenging projected future protein demands. Furthermore, faba bean’s high biological nitrogen fixation rates^8^ and long duration of nectar-rich, pollinator-friendly flowers^9^ provide important ecosystem services which mean that faba bean cultivation is increasingly seen as a key crop in sustainable intensification strategies. On the other hand, its partially allogamous mating system and estimated 13 Gb genome size, which puts it amongst the largest diploid genomes of any major crop, coupled with a low seed multiplication rate, have made it a challenging target for breeders^10^. Significant progress has been made in faba bean genomics and pre-breeding research. The mining of the first faba bean transcriptomes and development of SNP-based genetic maps, which showed strong collinearity with model legumes, set the scene for the identification of the WD40 transcription factor underlying the *Zero Tannin1* locus^11^ while a combination of high resolution mapping, transcriptomic, and metabolomic approaches led to the cloning of the *VC1* gene controlling seed content in vicine-convicine and paved the way for safer exploitation of the crop in the human food chain^12^. The lack of a reference genome sequence greatly complicated these studies, however, and improved faba bean genomic resources are urgently needed to accelerate crop improvement.

### The sequence of the giant faba bean genome

The 13 Gb faba bean genome (2n=2x=12) is one of the largest among diploid field crops (Extended Data Figure 1a, 1b) and its dominant repeat family members are longer^13, 14^ (up to 25 kb) than those in similarly sized polyploid cereal genomes^15^. The biggest of its six chromosomes holds the equivalent of an entire human genome. Although aiding cytogenetics^16^, these properties made genome assembly very challenging before the emergence of long and accurate reads. We sequenced the genome of the inbred line ‘Hedin/2’ with PacBio HiFi long reads to 20-fold coverage and assembled 11.9 Gb of sequence, more than half of which was represented by contigs longer than 2.7 Mb (Extended Data Table 1). Linkage information afforded by a genetic map (**Supplementary Table 1**) and chromosome conformation capture sequencing (Hi-C) data placed 11.2 Gb (94 %) into chromosomal pseudomolecules (Fig. 1a, Extended Figure 2a). Chromatin immunoprecipitation sequencing for centromeric histone H3 pinpointed the locations of the centromeres in the ‘Hedin/2’ assembly and arm ratios were consistent with karyotypes (Extended Data Fig. 2b). The single metacentric chromosome 1 was the only one to adopt a Rabl configuration, evident from the presence of both a main and an anti-diagonal on that chromosome in Hi-C interaction plots (Fig. 1a). This supports the notion that chromosome arms need to be of approximately equal size to spatially juxtapose in interphase. Some regions of the Hi-C contact matrices were empty for lack of mapped short reads (Fig. 1a). These white regions coincided with the location of enormous (up to 752 Mb) satellite arrays and aligned well with cytological maps of those repeats (Fig. 1 b-c). We also collected HiFi data (10-fold coverage) for the German variety ‘Tiffany’ and assembled these into a set of contigs with an N50 of 1.6 Mb and spanning 11.4 Gb (Extended Data Table 1). This level of completeness and contiguity was sufficient to arrange the contigs into pseudomolecules guided by the ‘Hedin/2’ reference (Extended Data Figure 3, Extended Data Table 1). In the future, the ‘Hedin/2’ assembly is expected to become the nucleus of a faba pan-genome.

**Figure 1.**
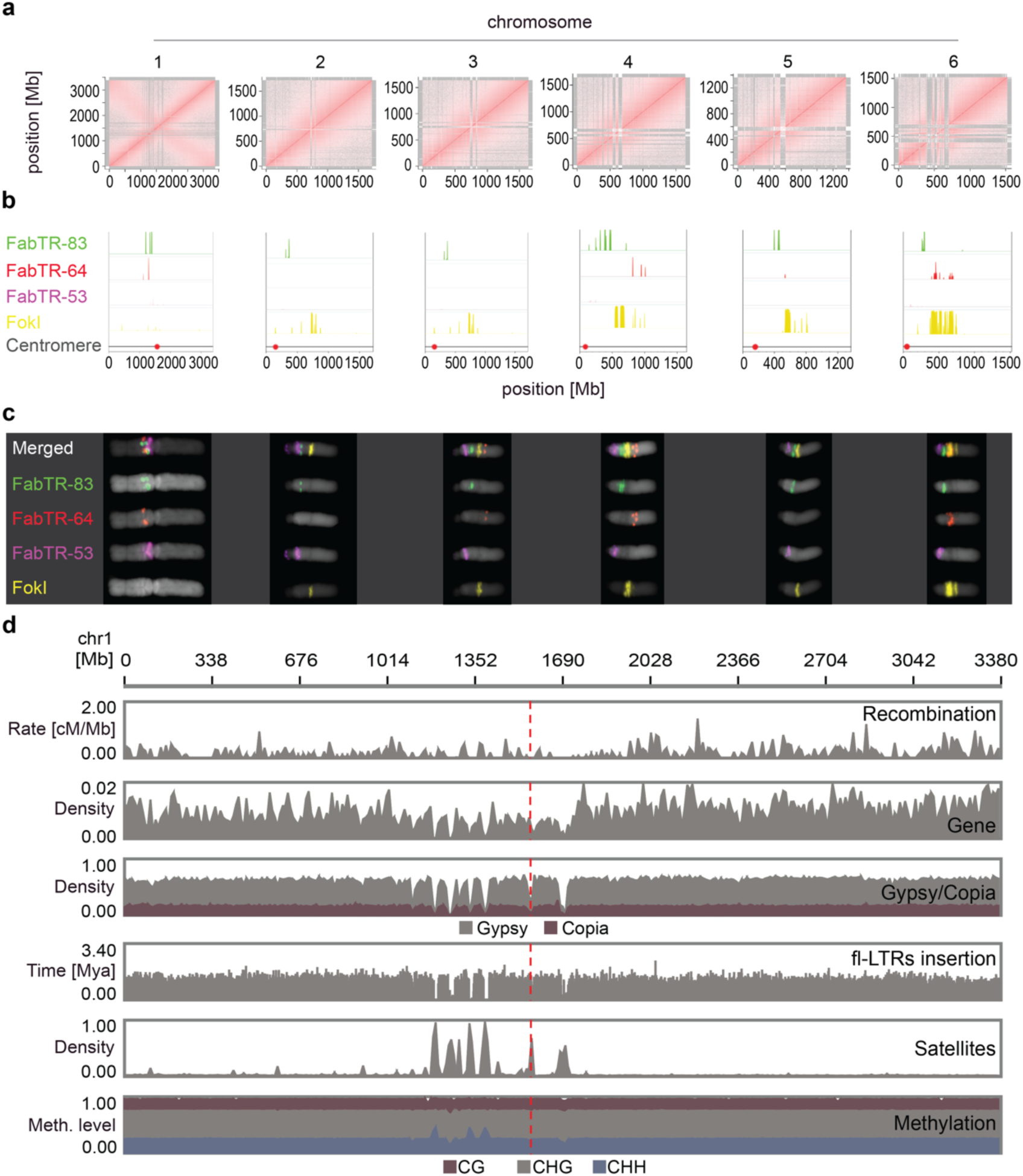
Gigabase-size chromosome-scale assembly of faba bean and its genomic features. **a,** Intrachromosomal contact matrix of assembled chromosomes. The red colour intensity indicates the normalized Omni-C Hi-C links between 1 Mb windows on each chromosome. The antidiagonal pattern in chromosome 1 represents the Rabl configuration. **b,** Distribution of major families of satellite repeats (FabTR-83, green; FabTR-64, red; FabTR-53, magenta; FokI, yellow) **c**, Distribution of major families of satellite repeats on metaphase chromosomes visualized by multi-colour Fluorescent *in situ* hybridization (FISH) (FabTR-83, green; FabTR-64, red; FabTR-53, magenta; FokI, yellow). **d**, Distribution of genomic components including recombination (cM/Mb), gene density, LTR retrotransposons of Gypsy and Copia, full-length LTR-RTs (fl-LTRs) insertion, satellite repeats and DNA methylation (CH, CHG and CHH context) on chromosome 1.

### Transposable elements, not polyploidy, have enlarged the genome

The genome sequence of ‘Hedin/2’ was annotated with RNA sequencing data from eight diverse tissues (**Supplementary Table 2**), resulting in a total of 34,221 protein-coding genes (Extended Data Table 2). A similar number of gene models (34,043) was also predicted in the ‘Tiffany’ assembly. The predicted ‘Hedin/2’ gene models captured 96% of single-copy orthologues conserved in Embryophyta according to the BUSCO metric (Extended Data Table 3). Gene density was uniform along the chromosomes (except for the positions of satellite DNA arrays) without the proximal-distal gradient typically observed for grass chromosomes^17^. Meiotic recombination displayed a similar distribution with an average of 27 genes per centimorgan (Fig. 1d, Extended Data Figure 4). Thus, despite its large genome, faba bean may be more amenable to genetic mapping than cereals, where up to a third of genes are locked in non-recombining pericentric regions^17^. Gene order was highly collinear and syntenic with other legumes (Fig. 2a). To further validate gene annotation, we aligned 262 *Medicago truncatula* genes related to symbiosis with rhizobia or arbuscular mycorrhizal fungi and found putative orthologues for them all. In addition, we verified using RNA-seq that a large subset of these genes were responsive to inoculation as expected ^18–20^ (**Supplementary Table 3**).

**Figure 2.**
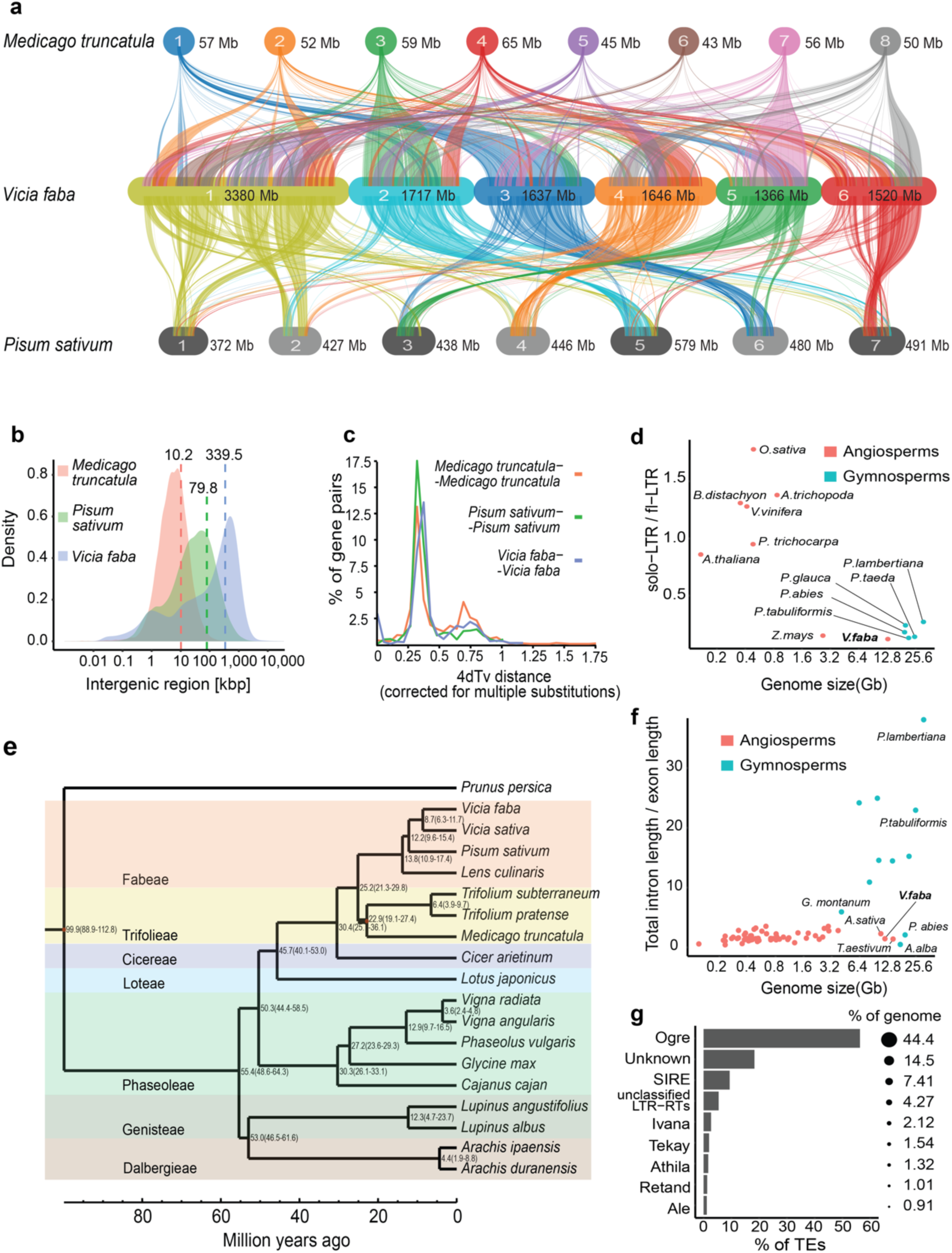
Evolution and synteny analysis in faba bean. **a**, Syntenic relationship of faba bean (middle) with Medicago (top) and pea (bottom). **b**, Comparison of intergenic length between legumes. **c,** Phylogenetic relationships between faba bean and other crop legumes in the Papilionoideae clade. The numbers on the branches indicate the estimated divergence time (million years ago, MYA). **d,** Distribution of the transversion rates at the four-fold degenerate sites (4dTv) of paralogous gene pairs of faba bean and other legumes. **e,** A plot shows the proportion of major retrotransposons **f,** Ratio of solo-LTR to full-length retrotransposons(fl-LTR) plotted against genome size in gymnosperms (green dots) and angiosperms (red dots). The ratio for other species was retrieved from Cossu *et al* (2017).

In contrast to gymnosperms with similarly gigantic genomes^21, 22^ introns in faba bean genes were not larger than in angiosperms with smaller genomes (Fig. 2f) but the intergenic space was much expanded (Fig. 2b). Moreover, the number of multi-copy gene families in faba bean was similar to related diploid species (Extended Data Figure 5), in contrast to soybean, which is considered a partially diploidized tetraploid^23^. Likewise, nucleotide substitution rates between paralogous and orthologous gene pairs place the last whole-genome duplication (WGD) event in the faba bean lineage at 55 million years ago (MYA) (Fig. 2c), well before the split from other Papilionoideae^24^ (Fig. 2e, Extended Data Figure 6), a taxon that also includes pea and lentil (*Lens culinaris*), species from which faba bean diverged around 12.2 and 13.8 MYA, respectively. Although we did not find evidence for a recent WGD in faba bean, more genes were duplicated in tandem than in pea and lentil (Extended Data Figure 7a). These duplications post-date the last WGD and occurred later than tandem duplications in *Arabidopsis thaliana* and *M. truncatula* (Extended Data Figure 7b), two species whose genomes were also rich in such events, and coincided with recent TE expansion. Overall, there were 1,108 syntenic clusters of tandemly duplicated genes in ‘Hedin/2’ and ‘Tiffany’, some of which differed in copy number. Notably, the agronomically relevant family of leghemoglobins had expanded (**Supplementary Table 5**). Despite this, the absence of a lineage-specific WGD or widespread gene family expansion means that the proliferation of repeat elements largely explains why the faba genome is more than seven times larger than that of its close relative common vetch (*V. sativa*).

Approximately 79% of the ‘Hedin/2’ assembly was annotated as transposon-derived (**Supplementary Table 6**). By far the largest group is the long-terminal repeat (LTR) retrotransposons (RLX), accounting for 63.7 % of the genome sequence. Other groups of TEs represent only minor fractions of the genome (**Supplementary Table 6**). Among the RLX, those of the *Gypsy* (RLG) superfamily outnumber *Copia* (RLC) elements by more than two to one (Fig. 1d, Extended Data Figure 4). The *Ogre* family of *Gypsy* elements alone make up almost half (44 %) of the genome, confirming its status as a major determinant of genome size in the Fabaceae^14^ (Fig. 2g). The great length of individual elements (up to 35 kb for *Ogre* and 32 kb of *SIRE*, the longest and second-longest elements), together with their abundance, partially explains the large size of the faba bean genome (Extended Data Figure 8). In addition, a large and diverse set of satellite repeat families that differ in their monomer sequences and genome abundance^25^ accounted for 9.4% of the total assembly length, with the most abundant satellite family *FokI* reaching 4% (0.475 Gb). *FokI*, together with several other highly amplified satellites, forms prominent heterochromatic bands on faba bean chromosomes (Fig. 1c). The TE density was remarkably invariable along all six chromosomes, mirroring gene density and recombination rate, and inverse to the density of satellite arrays (Fig. 1d, Extended Data Figure 4).

The persistence of retrotransposons as full-length copies can tell us about the balance between genome size expansion by retrotransposition and shrinkage by elimination through recombination. Modeling the solo-LTRs (sLTRs) as the product of the recombination between the LTRs of a single element, and assuming the canonical *Ogre* to comprise LTRs of 4,161 bp and in internal domain of 11,655 bp, the 395,657 sLTRs represent a loss of 6,26 Gb of DNA from the genome (55.6% of the current assembly size). This loss would be even greater if recombination between LTRs of different individual *Ogre* elements as well as DNA double-strand break (DSB) repair-mediated internal truncations, were considered. However, unlike plant species with smaller genomes, there were generally much fewer sLTRs in faba bean relative to the number of full-length LTRs, similar to large gymnosperm genomes (Fig. 2f), indicating slower removal than spreading of RLX^26^.

### Efficient genome-wide methylation

In addition to the relatively slow RLX elimination rate, it is also possible that lower levels of methylation could have accelerated TE proliferation through less efficient silencing. We found that most cytosines in the faba bean genomes were methylated: 95.8 % in CG, 88.2 % in CHG and 14% in CHH contexts, respectively (Fig. 1d, Extended Data Figure 4), placing it among the most highly methylated plant genomes^22^. Gene body methylation followed the canonical pattern (Fig. 3a) observed in other plants^27^: CG methylation was enriched in internal exons and introns (Extended Data Figure 9a), in contrast to low methylation in first exons, and may be related to transcriptional repression^28^. Genes with a high level of gene body methylation were more highly expressed in young leaf tissue (Extended Data Figure 9b) and also tended to be longer (average 3.3 kb). The elements of the major superfamilies of RLX, *Gypsy* and *Copia*, occupied 48% and 11% of the genome respectively, and were heavily methylated, more so in their bodies than their flanking regions (Fig. 3b). The most recent transposon burst occurred less than 1 MYA, but many structurally intact elements were between 3 and 5 million years old (Fig. 3c). Both young and old insertions were invariably methylated in all three sequence contexts (Fig. 3d). In contrast to other plant taxa^29^, RLX insertion times and methylation levels were uncoupled. Conspicuous islands of elevated CHH methylation also coincided with the abundant satellite repeat FabTR-83 (Fig. 3e), which accounts for 1.1% of the genome. Generally, the faba bean methylation machinery appeared fully functional, efficiently methylating all classes of repetitive elements, suggesting that methylation deficiency is unlikely to have played a role in genome expansion.

**Figure 3.**
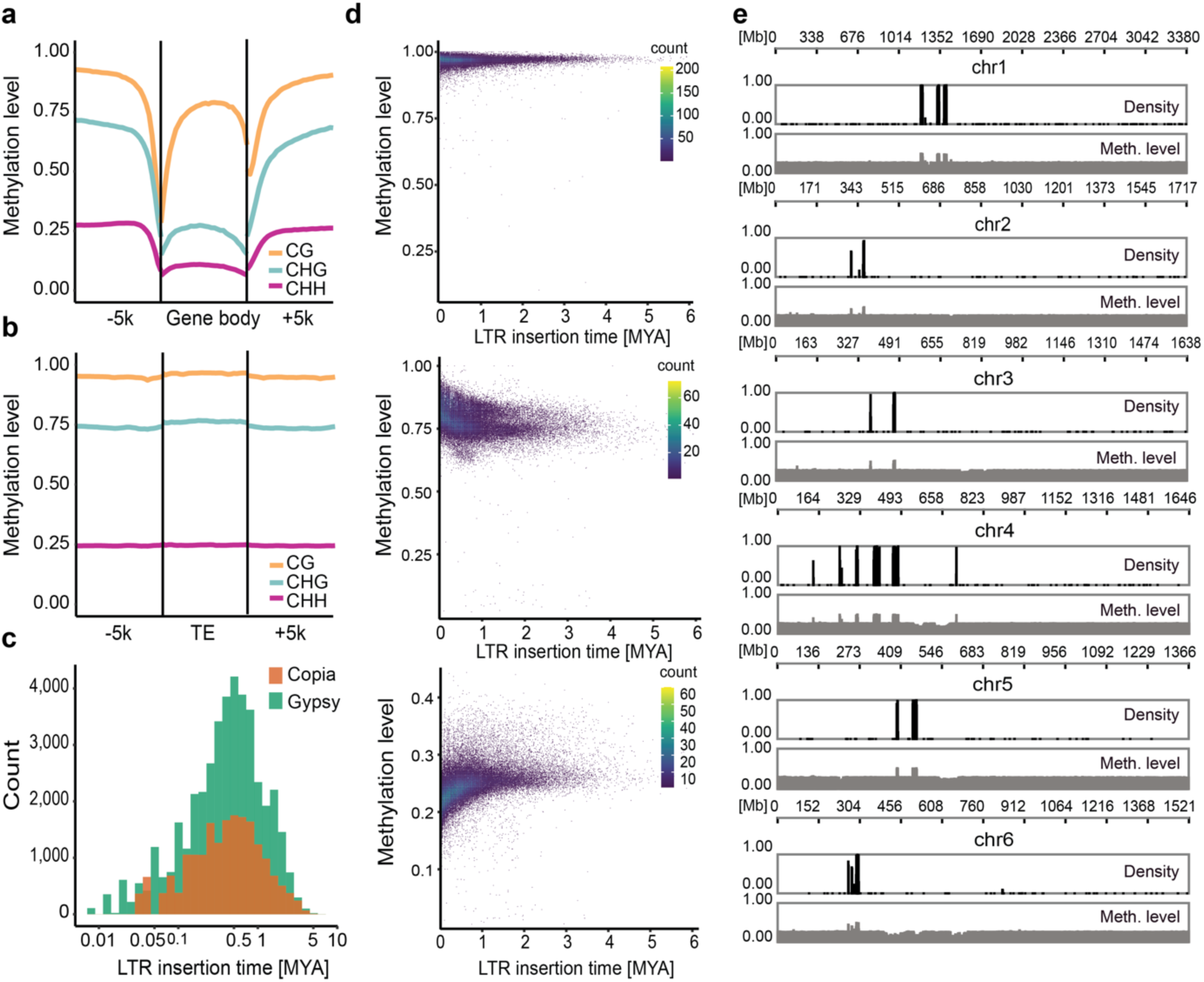
Methylation landscape in faba bean. **a**, Global distribution of DNA methylation levels at protein-coding genes including a 5 kb region upstream of the transcription start site (TSS) and downstream of the transcription end site (TES). **b,** DNA methylation patterns for TEs and their 5-kb flanking regions. **c,** Distribution of *Copia* (RLC)and *Gypsy* (RLG) retrotransposons based on insertion age. **d,** Landscape of CG (top), CHG (middle) and CHH (bottom) methylation with different TE insertion ages. **e,** CHH methylation peaks on FabTR-83 satellite repeats.

### Integration of QTL and variation data

The faba bean genome sequence provides a unified frame of reference for genetic mapping, gene expression profiling and comparative genomics. To assist the adoption of the new infrastructure among faba bean breeders and geneticists, we mapped markers from two commonly used genotyping platforms, the Illumina Infinium 1,536 single-nucleotide polymorphism (SNP) and the Illumina Oligo Pool Array (OPA) assays. Moreover, we projected genetic maps of both different bi-parental crosses and derived consensus genetic maps onto the genome assembly. This provided physical coordinates to quantitative trait loci (QTL) for disease resistance and phenology. Marker maps and QTL intervals can be browsed interactively at https://pulses.plantinformatics.io (Extended Data Figure 10).

The genome sequence has also paved the way for sequence-based genotyping. We mined the ‘Hedin/2’ assembly for oligonucleotide probes for use in Single Primer Enrichment Technology (SPET)^30^, a reduced-representation genotyping method with high-throughput and low per-sample costs. A panel of 197 accessions from a diversity collection were profiled with a 90,000 probe SPET assay with at least one probe in each predicted gene (**Supplementary Table 7**). Sequence reads were mapped to the ‘Hedin/2’ assembly and 1,081,031 segregating variants (SNPs) uniformly distributed along the genome were called, laying the foundations for high-resolution GWAS analysis (Fig. 1d, Extended Data Figure 11). Population structure analysis by model-based ancestry estimation and principal component analysis (PCA) divided the diversity panel into four groups, corresponding to their geographic origin (Extended Data Figure 12).

### Hilum colour mapping with candidate-gene resolution

To illustrate how a GWAS approach, in conjunction with multiple genome sequences, can help reveal the molecular basis for trait variation, we investigated the genetic control of seed hilum colour (Fig. 4a), which is an important quality trait^31^. We first carried out a GWAS for hilum colour using the diversity panel and identified a single prominent peak that was coincident both with the previously mapped *Hilum Colour (HC)* locus^31^ and peak homozygosity in a recessive pale hilum bulk of segregants from a cross between pale and dark hilum faba bean varieties (Fig. 4b). We found the most highly associated GWA marker in a polyphenol oxidase (PPO) gene residing in a cluster of eight fully intact and highly conserved *PPO* genes in the ‘Hedin/2’ assembly. In pea, PPO variation controls hilum colour, and a frameshifted, non-functional form of the single PPO copy residing at the syntenic *Pl* locus conferring a pale hilum is fixed in all modern pea varieties^32^. The pattern of pigmentation (Fig. 4a) and content in oligomeric phenolic compounds at the hilum surface in faba bean (Fig. 4c and Extended Data Figure 13) were very similar to those observed in pea^32^. Together with the genetic data, this indicates that differential PPO activity is responsible for hilum colour variation in both pea and faba bean, but it was unclear which faba bean PPO(s) may be causative.

**Figure 4.**
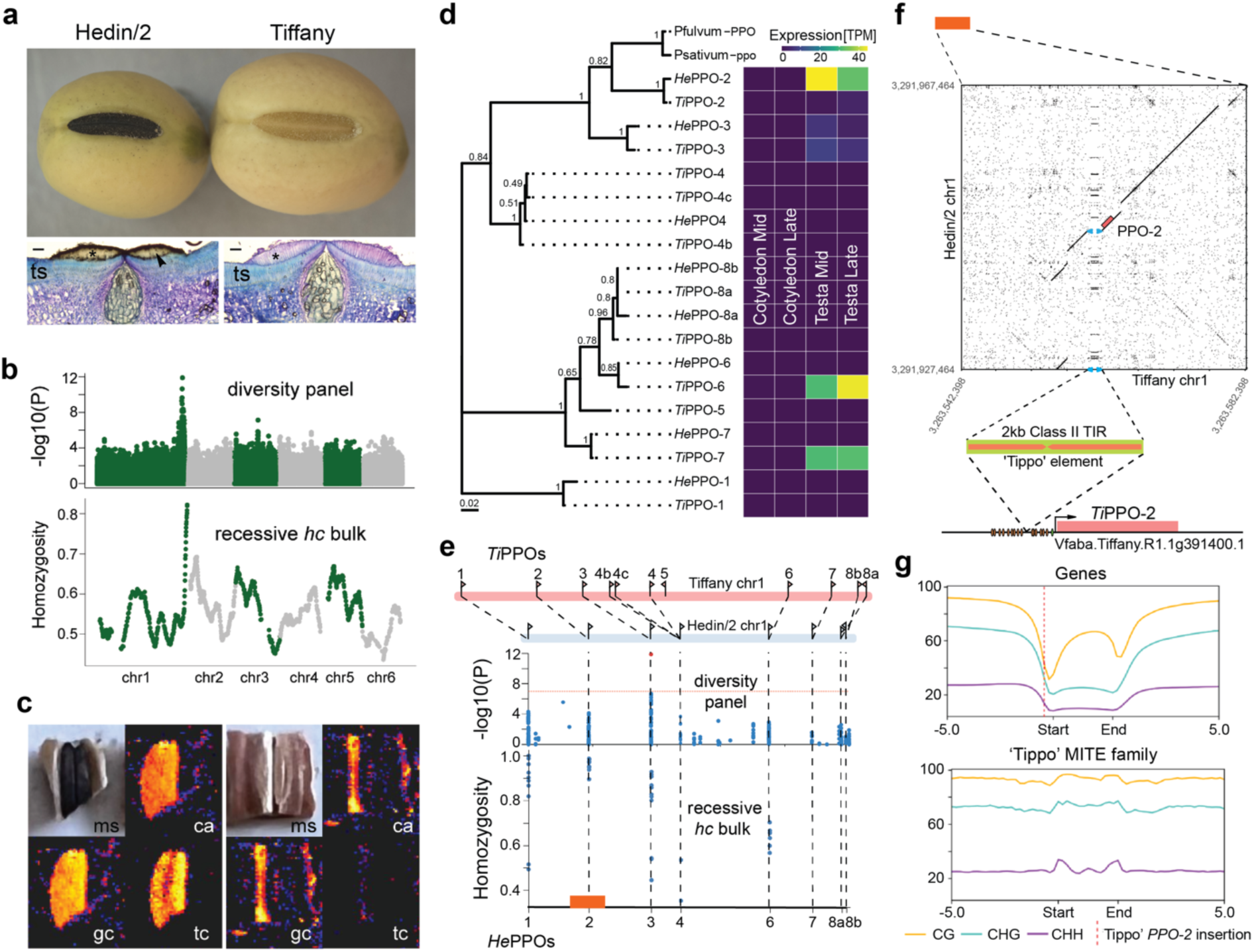
Rearrangements at the complex PPO locus give rise to changes in *PPO* expression and hilum colour. **a**, whole seeds of dark hilum Hedin/2 and pale hilum Tiffany are shown above light microscope images of a transverse section (ts) of the dark (left) and pale (right) hila where the scale bar is 20µm; **b**, genome-wide scans for association with hilum colour scored as a binary trait in the NORFAB diversity panel (top sub-panel) and for homozygosity of pale hilum parent alleles in an 84-component recessive pseudo-F2 pale hilum bulk; **c**, an optical image of Hedin/2 (left) and Tiffany (right) hilum specimens subjected to LDI/MSI (ms), and the LDI/MSI signal distribution for chlorogenic acid (ca), epi-gallocatechin (gc) and tetracosylcaffeate (tc) for the respective genotypes; **d**, phylogenetic tree showing the relationships between the causative pea gene and 8 and 11 PPO copies found in a tandem arrangement at the Hedin/2 and Tiffany *HC* locus respectively, with eight clades found forming the basis for a simplified PPO nomenclature and expression levels of each gene in testa tissue from mid and late pod filling expressed in Transcripts Per Million (TPM); **e**, from top to bottom: to-scale schematic of the chr1 PPO cluster showing order and orientation of PPOs in Tiffany and Hedin/2 with syntenic PPO copies are joined by dashed lines; closeup of hilum colour associations in the NORFAB diversity panel and homozygosity in the pale hilum (*hc*) bulk; **f**, dotplot detail of 20kb upstream and downstream of HePPO-2 and TiPPO-2 showing a c.2kb MITE, named ‘Tippo’ inserted in Tiffany amongst predicted transcription factor binding sites (brown ovals) in close proximity to the RNA Pol II binding site (TATA box – green oval) and transcriptional start site (arrow) of *PPO-2*; **g**, genome-wide methylation status of genes (top panel) compared to the Tippo MITE family (bottom panel).

To clarify, we compared the phylogeny and structure of the PPO clusters of the two fully sequenced genotypes ‘Hedin/2’ (dark) and ‘Tiffany’ (pale). *VfPPO-2* shared the highest level of identity with the causal pea gene *Psat1g2063360* (Fig. 4d), whereas the most strongly associated GWA marker was found in *VfPPO-3* and the pale hilum bulk homozygosity peak sat between *VfPPO-2* and *VfPPO-3*, suggesting that the causal polymorphism resided at the proximal end of the cluster (Fig. 4e). Structurally, apart from large differences in intergenic distances between syntenic PPOs caused mainly by *Ogre* insertions, the most striking features of the ‘Hedin/2’-‘Tiffany’ comparison were the triplication of *VfPPO-4* in ‘Tiffany’ and the absence of *VfPPO-5* in ‘Hedin/2’ (Fig. 4e), prompting us to investigate whether these structural variations were associated with variation in *PPO* gene expression. We first established that transcription of the *PPO* gene cluster was almost exclusively confined to the maternal testa tissue (which encompasses the hilum), rather than the cotyledon in both genotypes (Fig. 4d, Extended Data Figure 14, **Supplementary Table 8, Supplementary Table 9**). In ‘Hedin/2’ testa, *VfPPO-2*, and to a lesser extent *VfPPO-3*, accounted for nearly all *PPO* expression. In contrast, ‘Tiffany’ testa *PPO* expression was dominated by *VfPPO-6* and *VfPPO-7* (Fig. 4d). A detailed annotation and comparative repeat analysis of the PPO cluster region (Extended Data Figure 15) highlighted a c. 2kb AT-rich MITE insertion in the *TiPPO-2* promoter region (Figure 4f), which interrupts the sequence of predicted *VfPPO-2* promoter and belongs to a class of MITE associated with high levels of methylation (Figure 4g). Taken together, our results suggest that regulation of expression of *VfPPO-2* controls hilum colour variation in faba bean. Beyond suggesting a causative mechanism for pale hilum in faba bean, our analysis illustrates that increased copy number does not necessarily correlate with trait expression and emphasizes the utility of complete genome sequences from multiple genotypes.

## Discussion

Faba bean is one of the earliest domesticated crops. It was part of the Neolithic package of crops that the early farmers took with them as they left the Fertile Crescent^33^ and concern about the bean’s toxicity was voiced already in classical antiquity^34^. In the 21^st^ century, nutritional quality remains a central breeding goal: new faba bean varieties should be low in the alkaloid glycosides vicine and convicine as well as in tannins; essential amino acids should be balanced better to accommodate human dietary needs; seed phytate and protease inhibitors should be reduced to improve nutrient bioavailability; all while taking care not to compromise pest resistance and improving yield stability. Faba bean breeders can now face these challenges enabled by genomic resources and insights. Ubiquitous and frequent recombination will allow rapid introgression of new traits into elite material and permits powerful and broadly applicable mapping approaches exploiting the high SNP densities provided by SPET genotyping. Pinpointing causative variants can still be difficult in genomic regions with tandemly duplicated genes, but our investigation of hilum colour demonstrates that these challenges can be overcome using high-quality long-read assemblies coupled with transcriptomics. Repeats and their methylation influence genome evolution, but can also impact gene expression variation where the repeat elements insert within the regulatory regions of genes. Our rich genome-wide repeat annotation now sheds light on these effects, adding an important component to the genomics-based breeding platform. Expanding the platform further by cataloguing and exploiting as much of the segregating variation of domesticated faba bean as possible is now especially important, since we do not know the wild progenitor of faba bean. It appears to be an isolated species and does not hybridize with others in the genus *Vicia*^35^, effectively barring the use of wild relatives in faba bean breeding. Population-scale resequencing of mutants, genebank collections and elite cultivars along with pan-genome assemblies of representatives of major germplasm groups can now proceed supported by the resources and methods presented here.

## Acknowledgements

This research was supported by grants from the Innovation Fund Denmark (‘NORFAB’, 5158-00004B) to J.S. and FACCE-JPI ERA-NET SusCrop Profaba to S.U.A., D.M.O., F.S. and A.H.S.; German Leibniz Association in the frame of the Leibniz Junior Research groups (J118/2021/REPLACE) to M.J.; the German Federal Ministry of Education and Research (de.NBI, 031A536B) to H.G.; Novo Nordisk Fund (#NNF20OC0065157); the Jane and Aatos Erkko Foundation grant to A.H.S.; the Alexander von Humboldt Foundation in the framework of Sofja Kovalevskaja Award to A.A.G.; the Biotechnology and Biological Sciences Research Council award BB/P023509/1 to D.M.O.; the German Research Foundation (DFG) project number 497667402 to A.A.G. and B.U.; the Czech Science Foundation project number GACR 20-24252S to J.M.; the BMBF-funded de.NBI Cloud within the German Network for Bioinformatics Infrastructure (de.NBI) (031A532B, 031A533A, 031A533B, 031A534A, 031A535A, 031A537A, 031A537B, 031A537C, 031A537D, 031A538A). Parts of the public dataset curation for pulses.plantinformatics.io was funded by the Grains Research Development Corporation (GRDC). The authors wish to acknowledge CSC – IT Center for Science, Finland and ELIXIR CZ Research Infrastructure (Czech Ministry of Education, Youth and Sports grant no. LM2018131), for generous computational resources and thank Anne Fiebig for help with data submission.

## Author contributions

**M.J.** performed Hedin/2 genome assembly, methylation data analysis, hilum colour GWAS and wrote the first manuscript draft. **A.A.G.** performed Tiffany genome assembly and analysed seed coat and embryo expression data. **A.A.G, J.K. and B.I.** analysed synteny, tandem gene duplications and intron size variation. **J.K., A.A.G., and L.I.F** contributed with gene annotation and data submission. **S.U.A, D.M.O., L.I.F., D.A.** and **P.S.** designed the SPET assay. **D.A.** and **A.W.** performed hilum colour BSA mapping. **P.B.** carried out seed coat mass spectroscopy. **E.B.** and **L.L.J** contributed with SPET data analysis and submission. **P-E.C.** and **R.B.** generated and analyzed rhizobial and mycorrhizal symbiosis RNA-seq data. **K.C.** generated seed coat and embryo RNA-seq libraries. **J.D.** and **J.C.** estimated genome size using flow cytometry. **H.G., A.H.S., J.M., P.Ne., P.No., J.T.** and **J.K.** annotated and analysed repetitive elements. **A.Ha., P.T**., **B.U. and L.H.** performed gene functional annotation. **A.Hi., S.P.,** and **N.S.** generated methylome sequencing data. **S.K.** and **G.K-G.** contributed to PacBio sequencing, provided RNA-seq data used for variant discovery, and integrated and visualized QTL and variation data online. **A.K.** and **J.M** generated and analysed ChIP-seq data. **L.K.** and **P.S.** performed seed coat histochemistry. **P.K**. contributed seed coat analytical chemistry. **T.W.M.** carried out phenotype data validation. **M.M., D.M.O., A.H.S., S.U.A., I.S., G.A., M.N.**, and **H.K.** edited the manuscript with input from all authors. **J.O., L.K.N.**, and **A.S.** carried out SPET sample preparation. **L.K.N.** and **A.S.** carried out hilum colour phenotyping. **P.No.** developed the faba bean genome browser. **K.C.** and **T.R-S-H.** prepared seed coat and embryo RNA-Seq libraries. **L.A.R.** contributed cytogenetic analysis. **A.H.J.W.** and **R.S.** generated HiFi data. **H.Z.**: Gene family, evolution, diversity and phylogenetic analysis. **F.S.** provided plant material **S.U.A., J.S., N.T., A.M.T.** provided resources. **D.M.O.** and **D.A.** genotyped pure Hedin/2 stock and generated the genetic map. **S.U.A., A.H.S.** and **D.M.O.** and **M.J.** designed the study. **S.U.A.** coordinated the project.

## Additional information

Supplementary information is available for this paper.

Correspondence and requests for materials should be addressed to D.M.O, A.H.S, and S.U.A.

Data are available under European Nucleotide Archive (ENA) study ID PRJEB52541 and at www.fabagenome.dk.

## Online Methods

### Genome assembly

PacBio HiFi reads were assembled using hifiasm (v0.11-r302)^1^ with default parameters. The dovetail Omni-C data were aligned to the resulting contigs using minimap2^2^ to accurately order and orient the contigs. Similarly, the genetic markers from a consensus genetic map reported by Carrillo-Perdomo E et al^3^. and the 25K SNP array markers mapped in the NV644 x NV153 recombinant inbred lines (F6) were aligned to the preliminary contigs using minimap2 to assign contigs to chromosomes. Subsequently, the pseudomolecule construction was done with the TRITEX pipeline^4^. The final order and orientation of contigs in each chromosome were inspected and corrected manually with complementary support of Omni-C and NV644 x NV153 derived genetic map. The centromere regions were identified in each chromosome using ChIP-seq with the CENH3 (a centromere-specific histone H3 variant) antibody reported by Ávila Robledillo L et al^5^. Briefly, the raw reads from the ChiP-seq were trimmed by cutadapt (v.1.15)^6^ and mapped to the preliminary pseudomolecules using minimap2. The alignments were converted to BAM format using SAMtools^7^ and sorted by Novosort (V3.06.05) (http://www.novocraft.com). The read depth was then calculated in 100 kb windows. Finally, the order of each chromosome was determined with regard to centromere positions (short-to-long arm), matching with the karyotype map of faba bean.

### Estimation of genome size using flow cytometry

Nuclear genome size was estimated by flow cytometry as described previously^8^. Briefly, intact leaf tissues of the *V. faba* accession Hedin/2 and *Secale cereale* cv. Dankovske (2C = 16.19 pg DNA;^9^), which served as the internal reference standard, were chopped together in a glass Petri dish containing 500 μl Otto I solution (0.1M citric acid, 0.5% v/v Tween 20; Otto, 1990). The crude suspension was filtered through a 50 μm nylon mesh. Nuclei were then pelleted (300 × g, 2 min) and resuspended in 300 µl of Otto I solution. After 15 min of incubation on ice, 600 µl of Otto II solution supplemented with 50 µg/ml RNase and 50 µg/ml propidium iodide was added. Samples were analysed using a CyFlow Space flow cytometer (Sysmex Partec GmbH, Görlitz, Germany) equipped with a 532 nm green laser. The gain of the instrument was adjusted so that the peak representing G1 nuclei of the reference standard was positioned approximately on channel 100 on a histogram of relative propidium fluorescence intensity when using a 512-channel scale. Twelve Hedin/2 plants were sampled, and each sample was analysed three times, each time on a different day. A minimum of 5000 nuclei per sample were analysed and 2C DNA contents (in pg) were calculated from the means of the G1 peak positions by applying the formula: 2C nuclear DNA content = (sample G1 peak mean) × (standard 2C DNA content) / (standard G1 peak mean). The mean nuclear DNA content (2C) was then calculated for each species and DNA contents in pg were converted to the number of base pairs in bp using the conversion factor 1 pg DNA = 0.978 Gbp^10^.

### Genome size estimation and quality assessment

The distribution of the *k*-mer (*K*=101) frequency was estimated from PacBio HiFi reads using Jellyfish (v2.2.10)^11^. The output histograms were used to estimate the genome size and heterozygosity using findGSE^12^. The completeness of the assembly was assessed by two intendent approaches; i) self-alignment of HiFi reads to the assembly by minimap2 followed by SV calling using Sniffles^13^; ii) BUSCO (v3.0.2b)^14^ analysis with Embryophyta database 9.

### Enzymatic methylation sequencing

DNA for methylome sequencing was extracted using the Qiagen DNEasy Plant 96 kit in accordance with the manufacturer’s instructions, checked for intactness on a 1% agarose gel and quantitated using the Thermofisher Quant-iT™ PicoGreen™ dsDNA Assay. 200 ng of Hedin/2 genomic DNA was combined with 0.001 ng of CpG methylated pUC19 control DNA and 0.02 ng of unmethylated bacteriophage Lambda control DNA, then brought to a volume of 50 µl using EB buffer. The input DNA was sheared to 350-400 bp on the S220 Focused-ultrasonicator instrument (Covaris, Woburn, USA) using the following protocol: duty factor=10; peak incident power=175; cycles per burst=200; time=2 times 30 seconds. The sheared DNA was used to prepare a large insert NEBnext Enzymatic Methyl-seq library following the manufacturer’s instructions (https://www.neb.com/-/media/nebus/files/manuals/manuale7120.pdf). Four libraries were constructed with different sequencing indexes. Index PCR was performed with five PCR cycles to include indexes and amplify the libraries. The final libraries were quantified by qPCR, pooled at equimolar concentrations, and sequenced for 500 cycles (2 x 250 bp paired-end reads) on an SP-flow cell of the Novaseq6000 system (Illumina, Inc., San Diego, USA).

### Tiffany genome assembly

The distribution of k-mers (K=51) was estimated from PacBio HiFi reads using KAT (v2.4.2)^15^. The output histograms were used to estimate genome size and heterozygosity using findGSE. Assembly was performed using hifiasm v0.15.5-r350 (-l 0). The completeness of the assembly was assessed by aligning HiFi reads back to contigs and calling structural variants using cuteSV v1.0.11^16^. Despite there being no obvious heterozygous peak on the k-mer plots (Figure AS1), we observed a higher proportion of BUSCO duplicate genes in Tiffany compared to Hedin/2 and a slight over-estimation of genome size with findGSE. In addition, we also noted a number of short contigs with read coverage about half of the expected, suggesting the presence of regions of heterozygosity in the otherwise mostly homozygous genome. We therefore performed haplotig purging using purge_haplotigs v1.1.2^17^ (purge_haplotigs cov -l 3 -m 7 -h 25). Chromosome-level scaffolds were constructed with RagTag v2.0.1^18^ using the haplotig-purged assembly. To confirm the success of scaffolding, Hedin/2 and Tiffany chromosomes were aligned using GSAlign v1.0.22^19^. We compared two approaches for Tiffany annotation in order to choose one most suitable for comparative analyses: (i) individual annotation of genomes; (ii) a “transfer and gap fill” approach (Extended Data Figure 16), as described below. Overall, we observed that the transfer and gap fill approach resulted in more syntenic genes and more genes with the same CDS length. Visual inspection of annotation suggests that with the individual annotation approach, alternative transcripts of genes are annotated in some cases, which results in different CDS lengths for syntenic genes. The issue was largely resolved by using the transfer and gap fill approach.

### Gene model annotation

The repeat sequences were masked using RepeatMasker v4.1.1(http://www.repeatmasker.org) with a custom repeat library generated by RepeatModeler v2.0.1^20^ (using the Hedin/2 assembly). The gene annotation was conducted using BRAKER v2.1.6^21^ (etpmode, min_contig 10000). The RNAseq libraries (Table AS1) were aligned using STAR 2.7.8a^22, 23^. The protein database Viridiplantae OrthoDB v10.1^24^, merged with the translated sequences of the previously published *Vicia faba* transcriptome assembly^25^, was used as input for BRAKER, together with alignments generated by mapping the faba transcriptome assembly using GMAP v2020-10-14^26^. In addition, *Medicago truncatula* genes (“Mt4.0v2_Genes”) and *Pisum sativum* genes (“pissa.Cameor.gnm1.ann1.7SZR”) were aligned using GMAP v2020-10-14. The generated alignments were used to polish the BRAKER gene models. In order to account for any gene models missed by BRAKER prediction, but present in the Hedin/2 transcriptome assembly, the gene models from GMAP faba transcriptome alignments and BRAKER were compared using bedtools (2.30.0), retaining only the GMAP genes not having an intersection with the BRAKER gene models. For these genes, a further filtration was done to eliminate any short (<50 amino acids) translated proteins, in-frame stop codons, or low (< 200 reads) expression (featureCounts, subread 2.0.1^27^).

Completeness of the annotation was assessed for Hedin/2 and Tiffany by aligning one Iso-Seq dataset^28^ and assembled transcriptomes produced for cvs. Hiverna, Dozah and Farah. Transcriptomes were mapped using GMAP v2020-10-14 and comparisons between those mappings and the annotations were made using bedtools^29^. Gene models that had been removed by polishing, but which intersected mapped transcripts, were rescued if the transcript wasn’t a putative transposable element. *R* genes were detected on the unpolished and polished annotations using RGAugury^30^. *R* genes present in the unpolished annotation but not in the polished one were also rescued. The coding potential for each transcript was computed with CPC2^31^. The mRNAs with low coding potential were reclassified as lncRNAs. Genes of which at least 50% overlapped a transposable element domain were removed. Finally, any proteins that contained in-frame stop codons after phase correction were also removed. The completeness of the final gene set was assessed using BUSCOv5.2.2 with the embryophyta_odb10 and fabales_odb10 databases.

To ensure maximum comparability between annotations for Tiffany and Hedin/2 we employed the transfer and gap-fill strategy. In this approach, Hedin/2 complete protein coding gene annotations were transferred onto Tiffany assembly using Liftoff v1.6.1^32^. Transcripts with in-frame stop codons were removed and gaps between transferred genes were filled with Tiffany genes from the annotation obtained using the method described above, corresponding to genes removed due to in-frame stop codons and those which were Tiffany-specific.

### Symbiotic gene discovery

Total RNA sequencing was carried out for three biological replicates per condition. Eighteen libraries were prepared, and paired-end Illumina HiSeq mRNA sequencing (2×100bp RNA-Seq) was performed by GeneWiz (Leipzig, Germany), which produced around 2 × 70 million reads per library on average. Adaptor sequences were removed using CLC Genomics Workbench 11 (CLC Bio workbench, Qiagen, Aarhus, Denmark). Only inserts of at least 30 nt were conserved for further analysis. Reads were mapped to the Hedin/2 genome using the CLC Genomics Workbench 11 according to the manufacturer’s recommendations. The mapped reads for each transcript were normalized as total counts and used for calculating gene expression. Intact and broken pairs were counted as one. The total counts of each transcript under different conditions were compared using proportion-based test statistics^33^ implemented in the CLC genomic Workbench suite. This beta-binomial test compares the proportions of counts in a group of samples against those of another group of samples. Different weights were given to the samples, depending on their sizes (total counts). The weights were obtained by assuming a beta distribution on the proportions in a group, and estimating these, along with the proportion of a binomial distribution, by the method of moments. The result was a weighted *t*-type test statistic. We then calculated a false discovery rate correction for multiple-hypothesis tests^34^. Only genes showing a difference of 10 reads between compared conditions were considered as significantly expressed.

### Orthologous gene family identification

Genes from 19 legume species (Extended Data Table 4) were clustered to determine the orthologues relationship. The protein sequences from these species were aligned to each other using BLASTP v2.2.26^35^ with the parameter “evalue 1e-5”. The results were then used to cluster the gene families by OrthoMCL v2.0.9^36^.

### Phylogenetic analysis and divergence time estimation

The single-copy genes identified from 19 legume species (Extended Data Table 4) by OrthoMCL v2.0.9 were selected for the phylogenetic analysis. The four-fold degenerate synonymous site (4d locus) was extracted to build the evolutionary tree by PhyML^37^ and TreeBest (https://github.com/Ensembl/treebest). Molecular clocks and divergence times were estimated using MCMCTREE in the PAML v4.5^38^ package using the phylogenetic tree and the divergence time of known species (published literature or Timetree:http://www.timetree.org/).

### Whole genome duplication

The whole-genome duplication of *V. faba*, *M. truncatula* and *P. sativum* were estimated using the collinearity within each genome. First, synteny regions were identified using MCScanX ^39^. Then, the gene pairs in the synteny regions were used for 4dtv (four-fold degenerate transversions) calculation. The transversion rate was corrected by the HKY^40^ model. The synonymous (Ks) and non-synonymous (Ka) substitution was estimated by KaKs_Calculator 1.2^41^.

### Tandem duplicate gene discovery

Tandemly duplicated genes were also discovered using the CRBHits v0.0.4 package^42^ function tandemdups. To confirm the results, genes were also classified using DupGen_finder^43^, with *Arabidopsis thaliana* serving as the outgroup. *Vicia sativa* was excluded from TD analysis due to suspected fragmentation of its structural annotation, which could result in inflation in the number of genes annotated as TDs (**Supplementary Table 4**). The age of duplications was estimated using *T* = *K s* /2 *r*, *r =* 1.5 × 10 ^−8^. Ks was calculated using CRBHits using method ‘Li’. Synteny between Hedin/2 and Tiffany genes was analysed using CRBHits v0.0.4 package function rbh2dagchainer (type = "idx", gap_length = 1, max_dist_allowed = 20), which internally uses the DAGchainer algorithm^44^. Syntenic TDG clusters were discovered by connecting TDG clusters in individual genomes using the syntenic gene pairs found between Hedin/2 and Tiffany. To minimize the effect of unplaced genes on copy number variation (CNV) analysis, as unplaced genes can result in spurious CNV calls, we corroborated the synteny-based results with Orthofinder^45^ analysis. Only clusters that had the same or higher copy number in the same genotype, based on both synteny and Orthofinder results (for Orthofinder only genes on the matching chromosome and unplaced genes were considered), were retained for further analysis. Syntenic clusters were functionally annotated with Human Readable Descriptions (HRDs) using prot-scriber v0.1.0 (https://github.com/usadellab/prot-scriber).

### SPET library preparation and sequencing

Quantified genomic DNA using the Qubit 2.0 Fluorometer (Invitrogen, Carlsbad, CA) was used for library preparation, applying the Allegro Targeted Genotyping protocol (NuGEN Technologies, San Carlos, CA), which relies on a panel of probes. 20 ng/µL of DNA in solution was used as input following the manufacturer’s instructions. All libraries were quantified using the Qubit 2.0 Fluorometer and library size verified using the High Sensitivity DNA assay from Bioanalyzer (Agilent Technologies, Santa Clara, CA) or the High Sensitivity DNA assay from Caliper LabChip GX (Caliper Life Sciences, Alameda, CA). Libraries were also quantified by qPCR, using the CFX96 Touch Real-Time PCR Detection System (Bio-Rad Laboratories, Hercules, CA). Samples were sequenced at IGA Technology Services (IGATech, Udine, Italy). DNA sequencing was performed on the Illumina NovaSeq 6000 (Illumina, San Carlos, CA) in a 2×150 PE configuration, generating an average of 7.73 M sequenced read pairs per accession.

### Hilum colour and Histology

To examine hilum morphology, seed coat containing hilum from inbred lines Hedin/2 (dark hilum) and Tiffany (pale hilum) were dissected from mature dry seed, saturated with 2% sucrose solution under vacuum for 1 hour and embedded in cryo-gel media (Cryo-gel Leica). Samples were cut in cryotome (Leica CM1950, US) into 15µm transversal sections) and stained with Toluidine blue O (0.01 %, w/v in water; Sigma Aldrich, CZ) as previously described^46, 47^. Observation and photography was done on an Olympus BX 51 microscope (Olympus Corp., Tokyo, Japan) in bright field and figures were documented with an Apogee U4000 digital camera (Apogee Imaging Systems, Inc., Roseville, CA, USA). For the investigation of metabolite content of surface layers of the hilum by laser desorption-ionization imaging mass spectrometry (LDI-MS), seeds were mechanically cracked and hila with small part of surrounding tissue were separated from the rest of seed coats using microscissors (MicroSupport, Shizuoka, Japan), fixed using a double-sided tape on MALDI plates with outer surfaces facing up and analysed as previously described^46, 48^. LDI-MSI experiments were done using a high-resolution tandem mass spectrometer (HRTMS) Synapt G2-S (Waters, Milford, USA). The vacuum MALDI ion source used was equipped with a 350 nm 1 kHz Nd:YAG solid state laser. Parameters of the mass spectrometer were set as follows: Extraction voltage at 10 V, Collision energies: Trap collision energy (TrapCE) 4 eV, Transfer collision energy (TransferCE) 2 eV. TrapCE at 25 eV and LM resolution at 10 were used for MS/MS experiments. Instrument calibration was done using red phosphorus (1 mg.mL^-1^, suspension in acetone). Mass imaging data collection was driven by HDImaging 1.5 software (Waters). The laser beam size was 60 μm. Spectra were collected in positive and negative ionization mode with laser energy 300 arb. Laser repetition rate was set up at 1000 Hz. Mass range was 50–1200 Da.In order to fine-map the *HC* locus, a cross was made between inbred lines Disco (♀pale) and Hedin/2 (♂dark). F4 seed from 337 F3 progeny of 21 F2 individuals shown by flanking marker analysis to be heterozygous across the *HC* interval were scored for hilum colour resulting in a 253 dark:84 pale hilum ratio (χ^2^ = 0.00098, p-value – 0.9749 for fit to expected 3:1 ratio). A pool composed of equimolar quantities of DNA from each of the 84 recessive pseudo-F2 individuals was created and subjected to SPET re-sequencing alongside DNA samples of the parent lines. In order to study expression of the PPO gene family in mid-to late-pod fill, individual plants of Hedin/2 and Tiffany, were grown in the glasshouse until the most mature pods on lower nodes had almost reached maturity and the uppermost nodes were still in flower, giving a gradient of seed development. All pods were then harvested and dissected into pod wall, testa, cotyledon, funicle and embryo axis samples (Extended Data Figure 17); fresh weights of each tissue recorded. Because all pods on a given node are not fertilized synchronously nor necessarily progress through development at the same rate, and based on insights from our prior studies of faba bean seed development^49^, we categorised individual pods into mid- and late-pod-fill stages in terms of the ratio of cotyledon weight to the total seed weight (Extended Data Figure 18).

### Comparative Sequence Analysis

To identify *PPO* homologues in Hedin/2 and Tiffany proteomes, the protein sequence from the pea *PPO1/Pl* gene (Psat1g206360) was used as a BLAST query. Multiple sequence alignment of PPO protein sequences was performed using Clustal Omega v1.2.4. The evolutionary history was inferred using the Maximum Likelihood method and JTT matrix-based model as implemented in MEGA X with 100 bootstrap replicates. The complete *PPO* regions (from the beginning of the first to the end of the last PPO gene and 10,000 bp flanking sequences on both sides) were extracted and aligned using minimap2 v 2.24-r1122. Then, 20,000 bp downstream and upstream from the transcription start of *PPO-2* were extracted and similarity between sequences was visualised using FlexiDot.

### Gene Expression Analysis

RNA was extracted from 100mg of flash-frozen dissected tissue (testa and cotyledon) using a Sigma Spectrum Kit (STRN250) according to the manufacturer’s directions, except that incubation was made at room temperature after DNA digestion. While extraction of RNA from cotyledons was performed exactly as per manufacturer’s specifications, testa tissue was disrupted in an extraction buffer consisting of CTAB, PVP, 2M Tris pH 8, 0.5M EDTA pH 8, 4M NaCl, spermidine and beta mercaptoethanol, followed by precipitation with 8M lithium chloride (instead of the kit’s lysis step). Total RNA was quantified using Qubit RNA IQ assay and normalised prior to preparation of directional mRNA sequencing libraries using standard methods. Between 4.1 and 5.6 million Illumina PE150 short reads per library (3x replicates, 2x tissues, 2x genotypes) were generated. Hedin/2 and Tiffany gene expression was quantified using Kallisto v 0.44.0 by pseudo-aligning RNA-Seq reads to respective reference transcripts. Transcript level abundance was converted to gene level abundance using tximport.

## Extended Data Tables

**Extended data Table 1.**
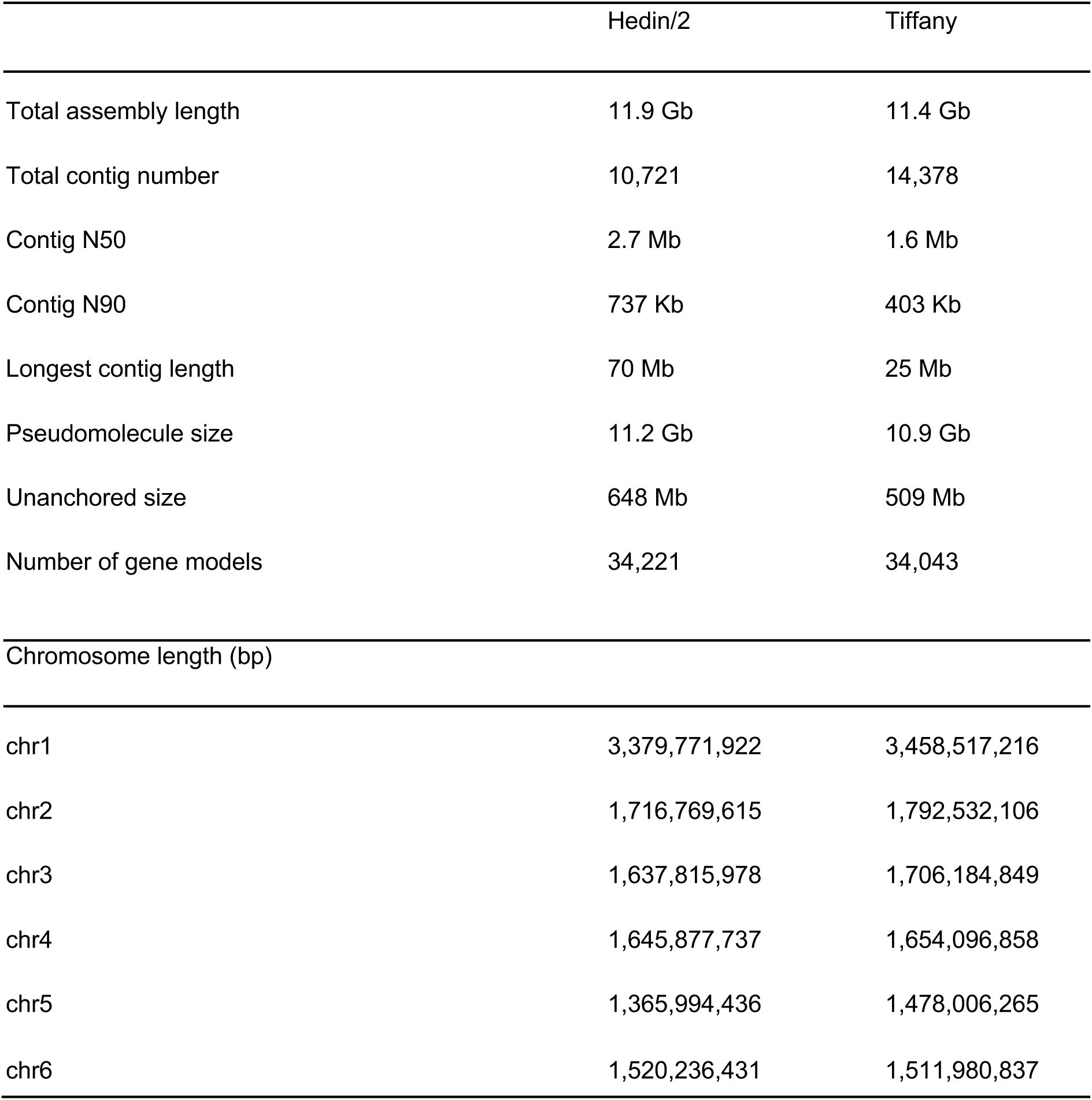
Summary of Genome assembly statistics.

**Extended data Table 2.**
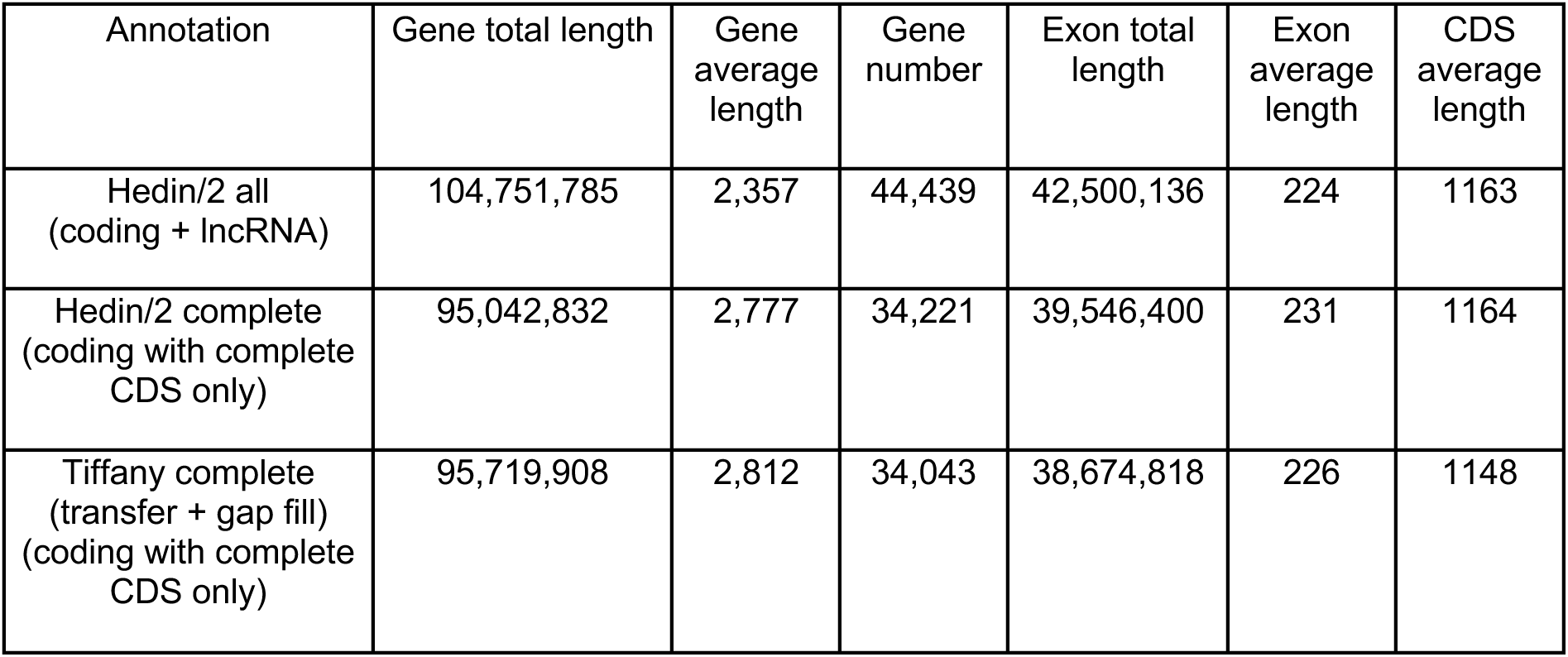
Structural annotation statistics for Hedin/2 and Tiffany assemblies.

**Extended data Table 3.** Annotation BUSCO completeness.

| **Annotation** | **Complete [S:Single copy, D:Duplicated]** | **Fragmented** | **Missing** | **Groups checked** | **BUSCO database** |
| --- | --- | --- | --- | --- | --- |
| Hedin all | C:96.8%[S:78.0%,D:18.8%] | 2.0% | 1.2% | 255 | embryophyta_odb10 |
|  | C:94.7%[S:90.6%,D:4.1%] | 0.9% | 4.4% | 5366 | fabales_odb10 |
| Hedin/2 complete | C:96.8%[S:78.0%,D:18.8%] | 2.0% | 1.2% | 255 | embryophyta_odb10 |
|  | C:94.5%[S:90.4%,D:4.1%] | 0.9% | 4.6% | 5366 | fabales_odb10 |
| Tiffany complete (transfer+gap fill) | C:94.9%[S:77.3%,D:17.6%] | 2.7% | 2.4% | 255 | embryophyta_odb10 |
|  | C:90.2%[S:85.7%,D:4.5%] | 1.0% | 8.8% | 5366 | fabales_odb10 |

**Extended data Table 4.** Summary statistics of gene families.

| **Species** | Gene number | Genes in families | Un-clustered genes | Family number | Unique families | Average genes per family |
| --- | --- | --- | --- | --- | --- | --- |
| *Arachis duranensis* | 32,899 | 27,703 | 5,196 | 17,302 | 341 | 1.6 |
| *Arachis ipaensis* | 35,785 | 28,950 | 6,835 | 17,438 | 573 | 1.66 |
| *Cajanus cajan* | 30,842 | 26,425 | 4,417 | 16,507 | 256 | 1.6 |
| *Cicer arietinum* | 28,260 | 25,996 | 2,264 | 15,181 | 322 | 1.71 |
| *Glycine max* | 51,431 | 42,082 | 9,349 | 17,633 | 396 | 2.39 |
| *Lens culinaris* | 38,624 | 33,149 | 5,475 | 16,672 | 929 | 1.99 |
| *Lotus japonicus* | 25,460 | 23,104 | 2,356 | 14,613 | 324 | 1.58 |
| *Lupinus albus* | 37,837 | 30,588 | 7,249 | 17,318 | 289 | 1.77 |
| *Lupinus angustifolius* | 33,663 | 31,447 | 2,216 | 16,644 | 200 | 1.89 |
| *Medicago truncatula* | 31,571 | 28,978 | 2,593 | 16,857 | 368 | 1.72 |
| *Phaseolus vulgaris* | 27,263 | 25,366 | 1,897 | 16,890 | 112 | 1.5 |
| *Pisum sativum* | 43,677 | 32,990 | 10,687 | 18,684 | 1,476 | 1.77 |
| *Prunus persica* * | 26,631 | 22,127 | 4,504 | 14,490 | 604 | 1.53 |
| *Trifolium pratense* | 33,342 | 31,284 | 2,058 | 17,108 | 388 | 1.83 |
| *Trifolium subterraneum* | 39,578 | 29,312 | 10,266 | 17,066 | 740 | 1.72 |
| *Vicia faba* | 34,221 | 28,970 | 5,211 | 17,721 | 700 | 1.63 |
| *Vicia sativa* | 43,831 | 34,301 | 9,530 | 18,275 | 1,246 | 1.88 |
| *Vigna angularis* | 26,445 | 24,753 | 1,692 | 16,506 | 111 | 1.5 |
| *Vigna radiata* | 26,715 | 24,917 | 1,798 | 16,511 | 144 | 1.51 |

\*outgroup.

## Extended Data Figures

**Extended Data Figure 1.**
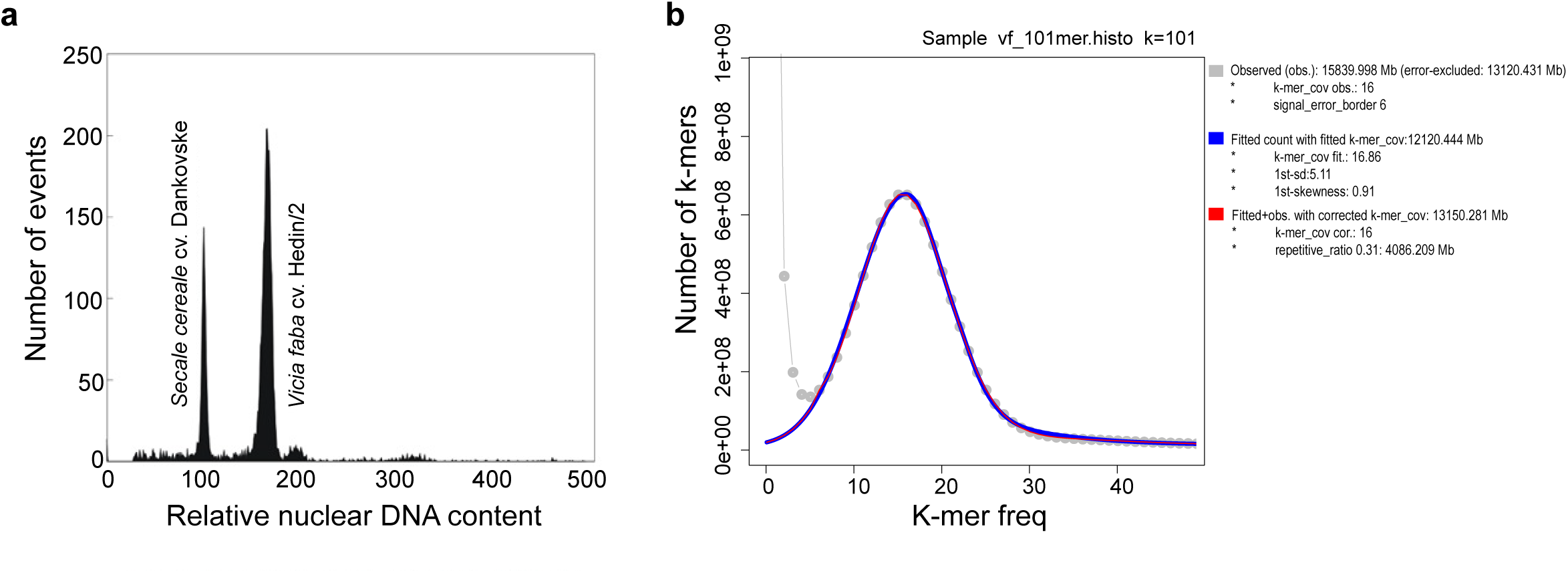
K-mer and flow-cytometry based estimation of faba bean cv. Hedin/2 genome size. a, Histogram of relative DNA content obtained using flow cytometric analysis of fluorescence of cell nuclei stained by propidium iodide. The nuclei were isolated simultaneously from leaf tissues of V. faba cv. Hedin/2 and Secale cereale cv. Dankovske, which served as an internal reference standard. The ratio of peaks representing G1 nuclei informs the ratio of genome sizes. b, k-mer based estimation of genome using Hedin/2 raw HiFi sequencing data.

**Extended Data Figure 2.**
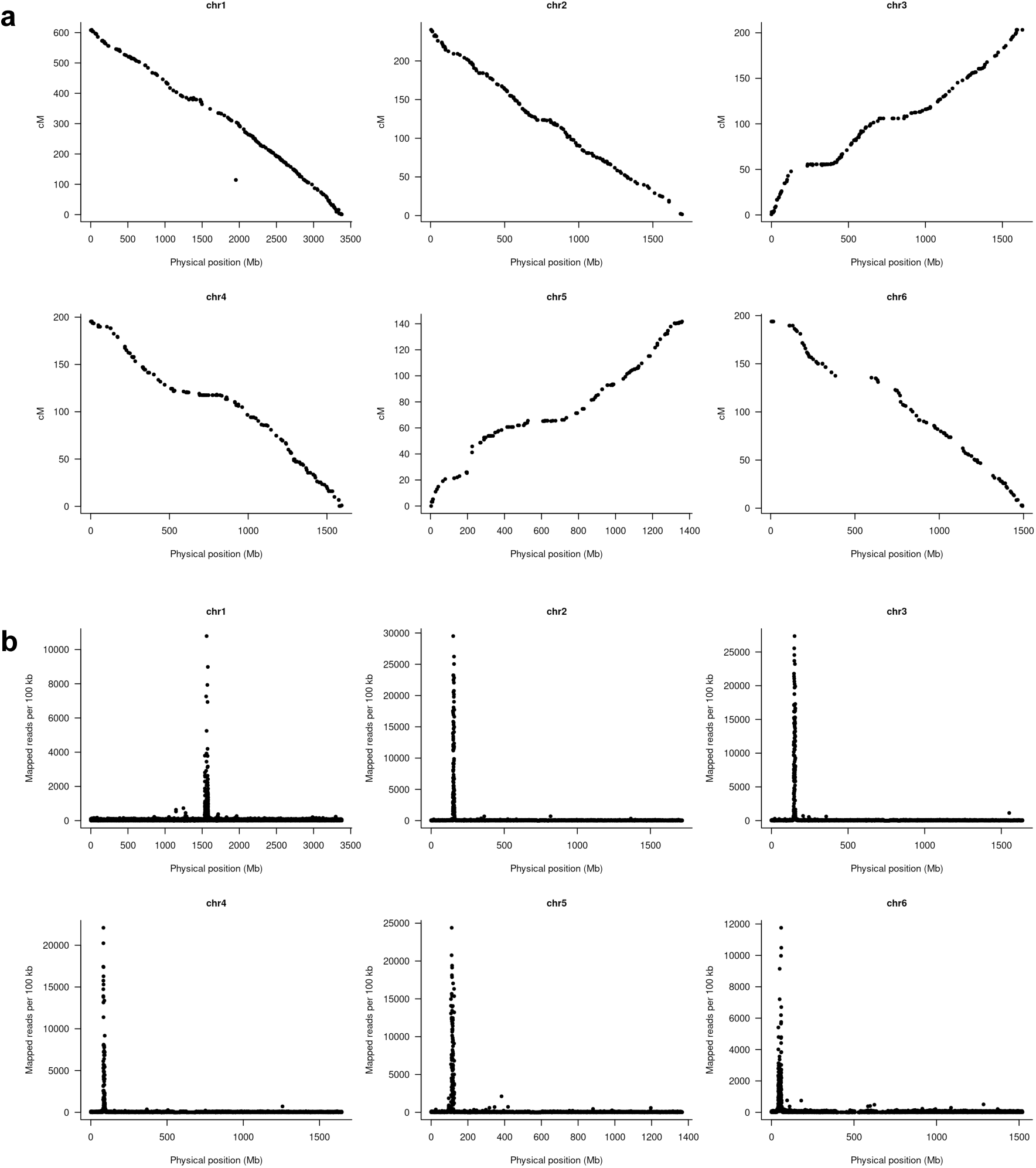
Collinearity of physical and genetic maps and ChIP-seq localisation of centromeres. a, Collinearity of physical and genetic maps. The antidiagonal alignments in chr1, chr2, chr4, and chr6 were a result of the arbitrary orientation of linkage groups in prior genetic maps. b, chromatin immunoprecipitation sequencing (ChIP-seq) localization of centromeres.

**Extended Data Figure 3.**
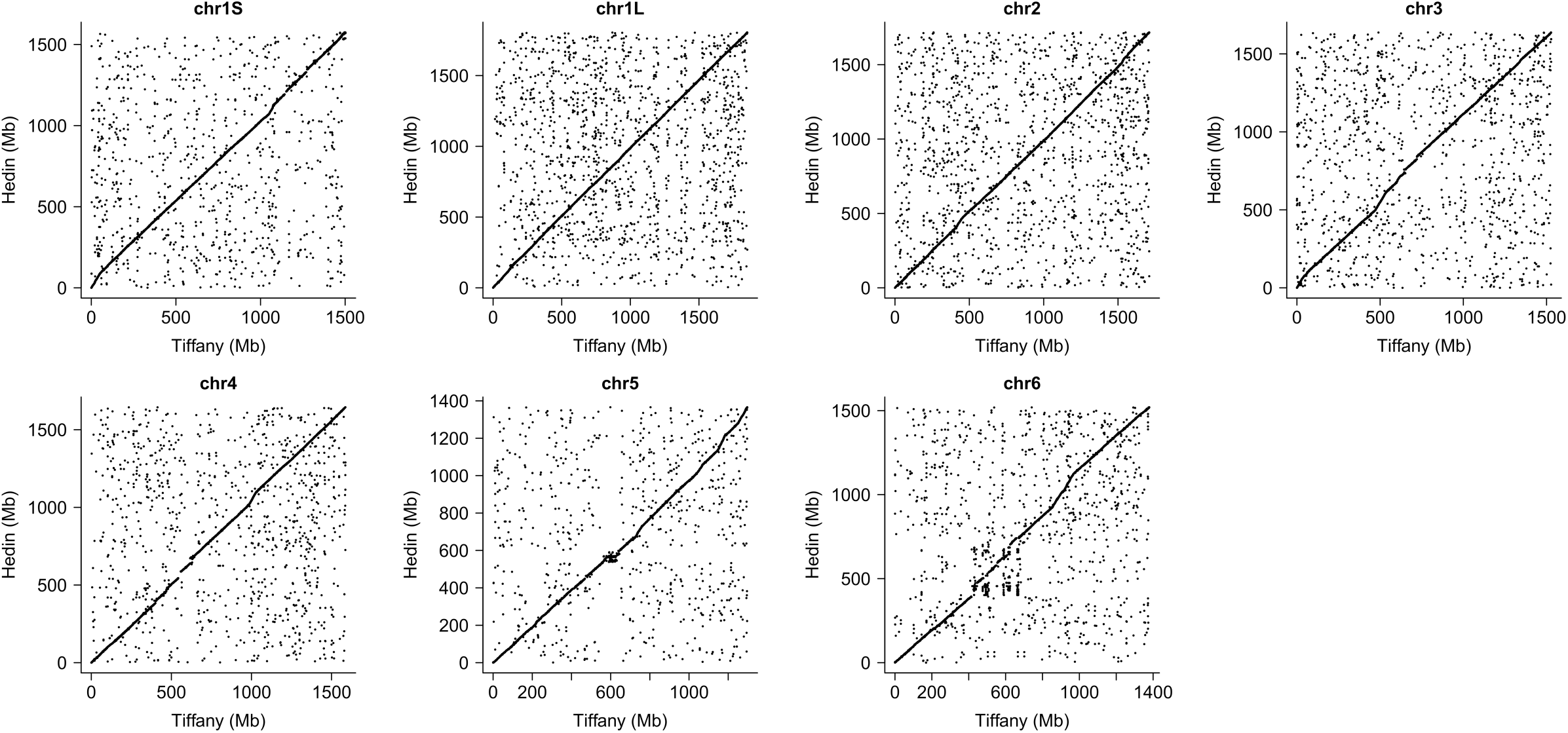
A dot plot showing whole genome alignment between Hedin/2 and Tiffany.

**Extended Data Figure 4.**
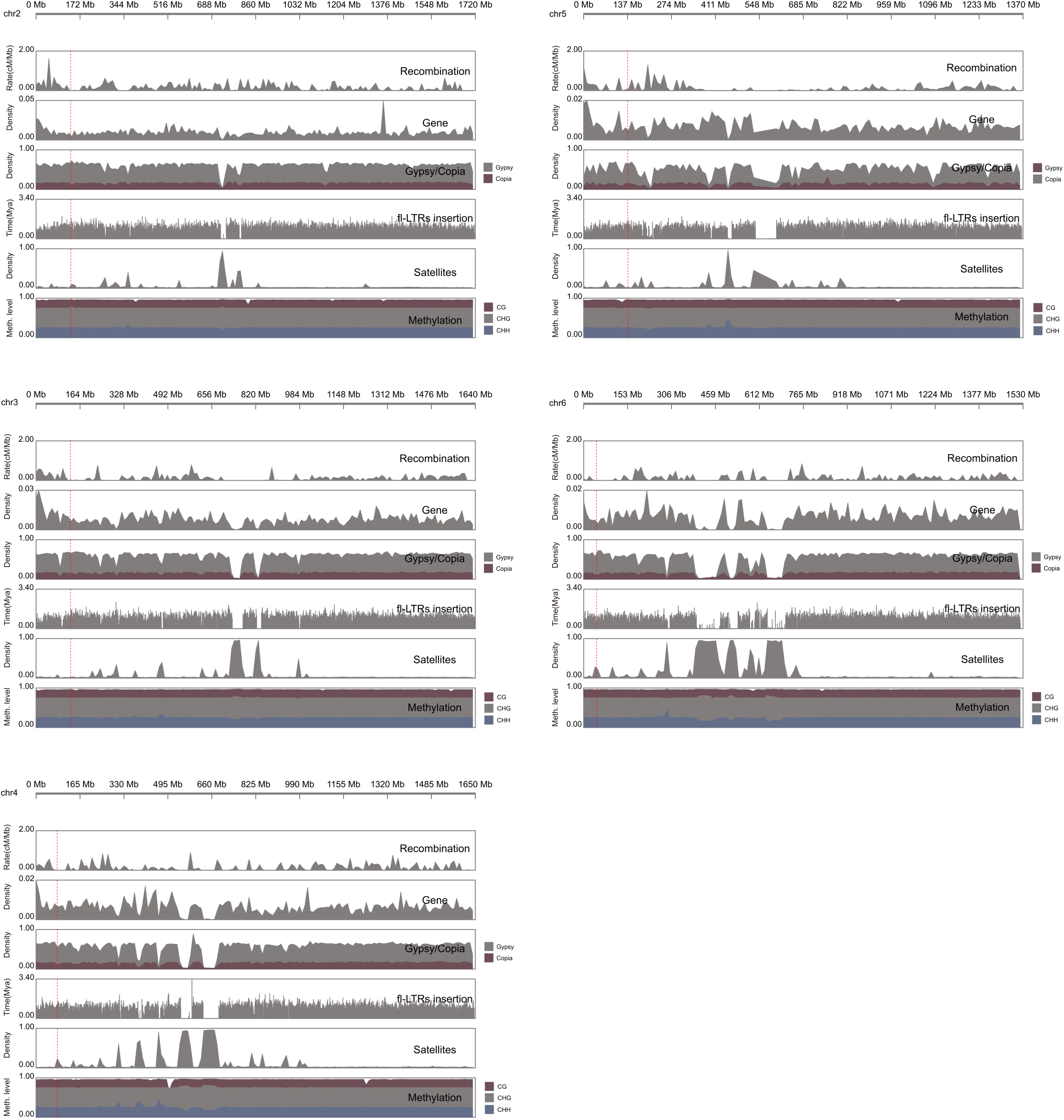
Distribution of genomic components including recombination (cM/ Mb), gene density, retrotransposons of Gypsy and Copia superfamiles, full-length (FL) insertions, satellite repeats and DNA methylation (CH, CHG and CHH context) on chromosome 2 to 6.

**Extended Data Figure 5.**
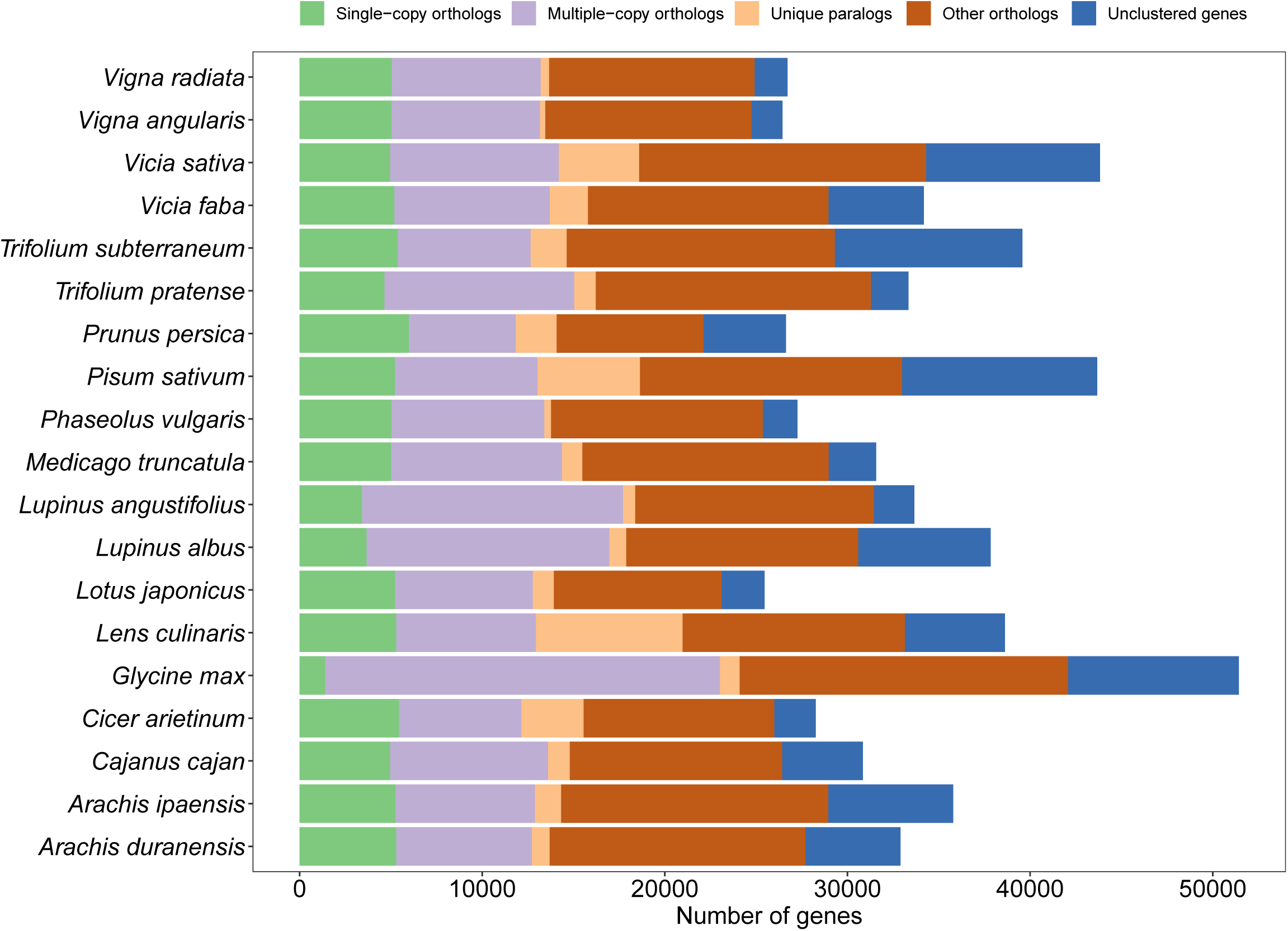
A plot of homologous gene number in each of 19 species. Single-copy orthologues, single-copy homologous genes in the gene families shared among species; multiple-copy orthologues, multicopy homologous genes in gene families shared among species; unique paralogues, genes of the strain unique to the family; other orthologs, all other genes; unclustered genes, genes not clustered into any family The horizontal bars represent the number of protein-coding genes.

**Extended Data Figure 6.**
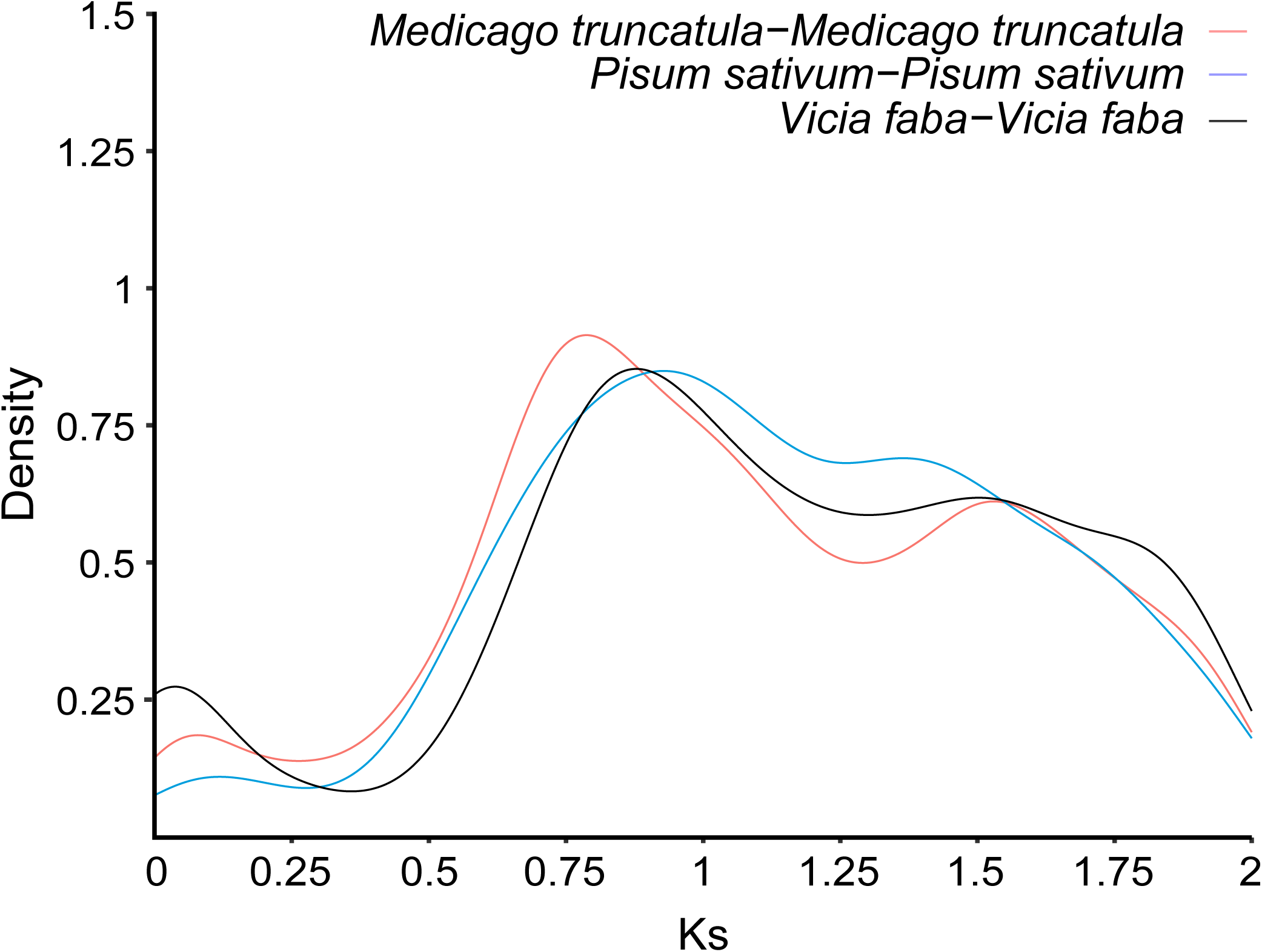
Distribution of the synonymous substitution rate (Ks) between paralogous gene pairs of faba bean and other legumes.

**Extended Data Figure 7.**
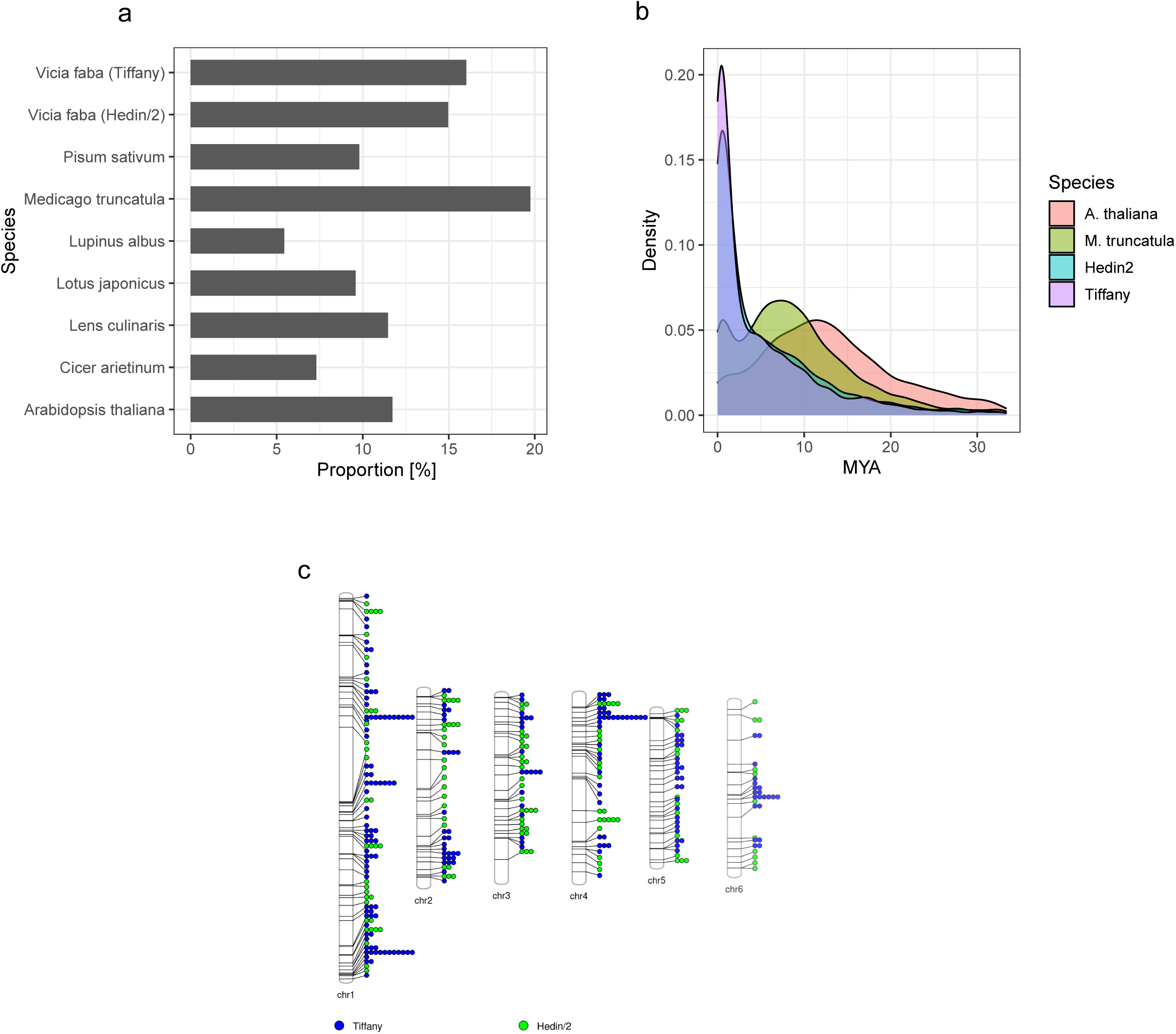
**a,** Proportion of genes tandemly duplicated. **b,** Estimated age of tandem duplications. **c,** Tandem duplicate gene copy number variation between Hedin/2 and Tiffany. Number of circles - difference in copy number (number extra copies in one of the genotypes). Colour – genome, which carries more copies.

**Extended Data Figure 8.**
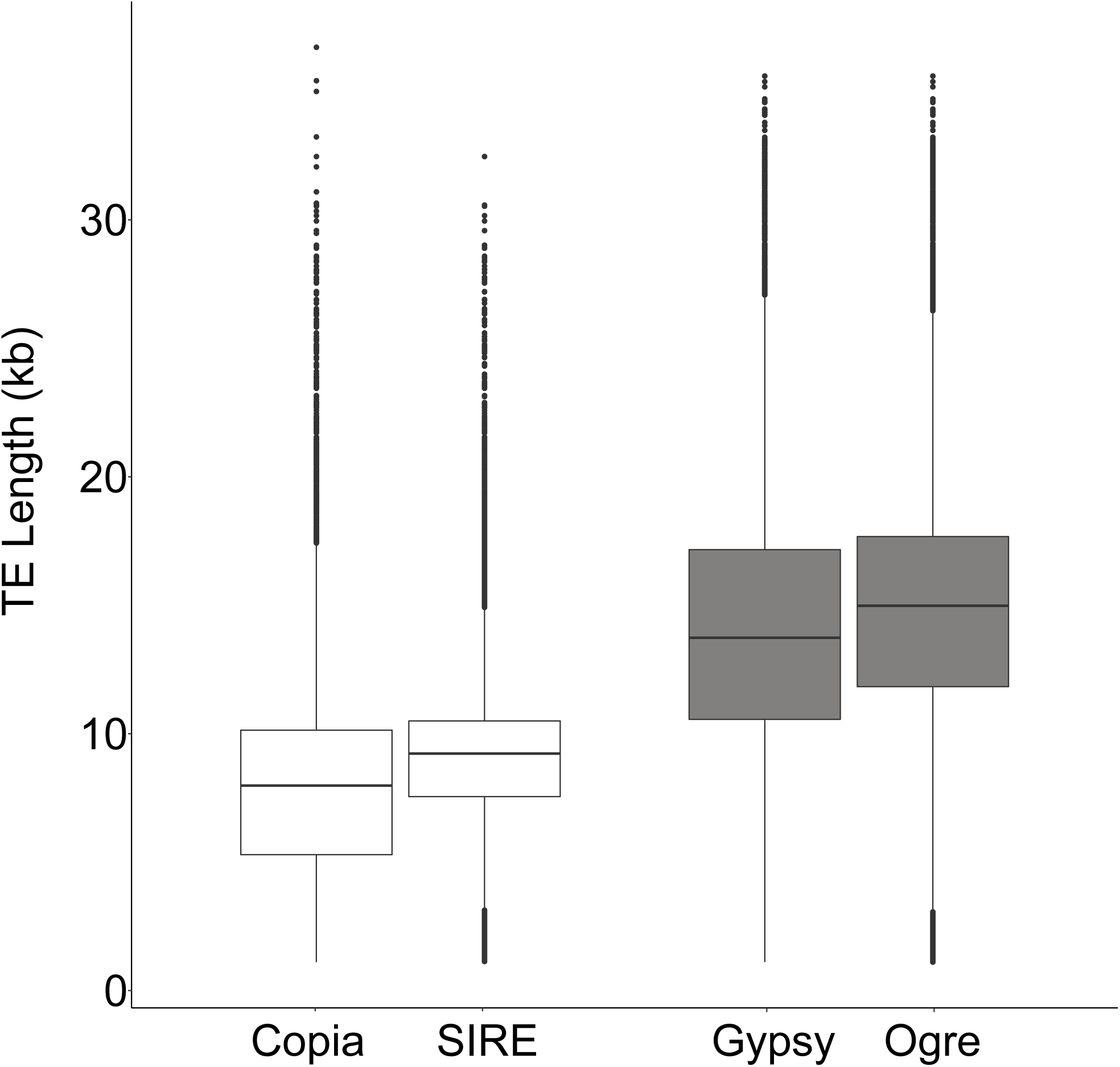
Full-length representation of Copia and Gypsy super family. Ogre and SIRE are the largest size element in Gypsy and Copia respectively.

**Extended Data Figure 9.**
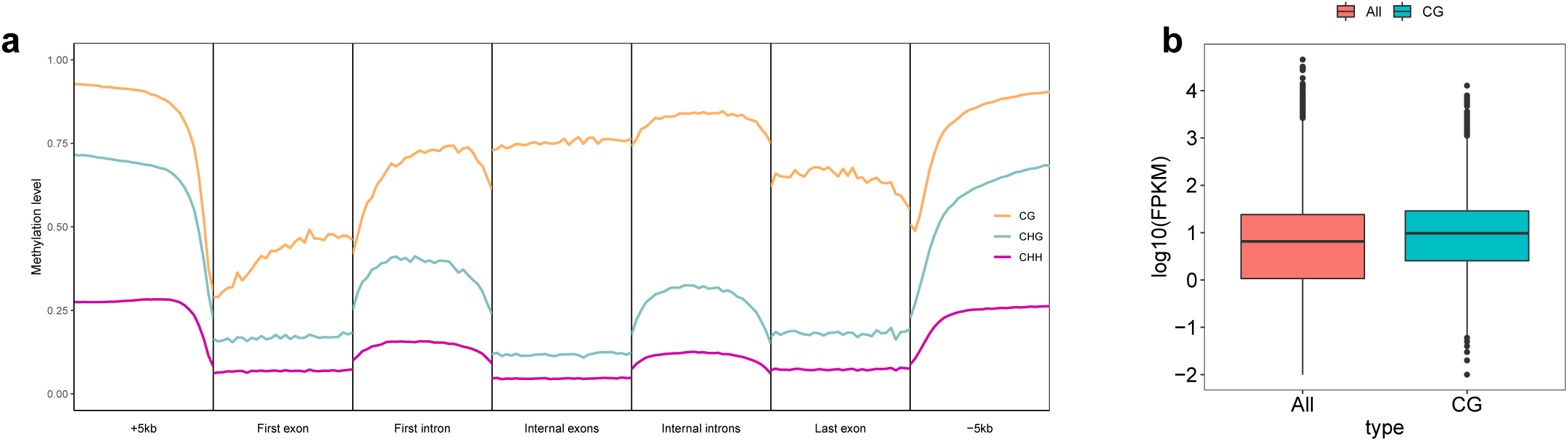
**a,** Methylation pattern based on gene features including first exon, first intron and last exon, last intron together with 5-kb flanking regions in Hedin/2 genome. **b,** Expression comparison between all annotated genes and gene-body methylated genes (CG context) in young leaf tissue.

**Extended Data Figure 10.**
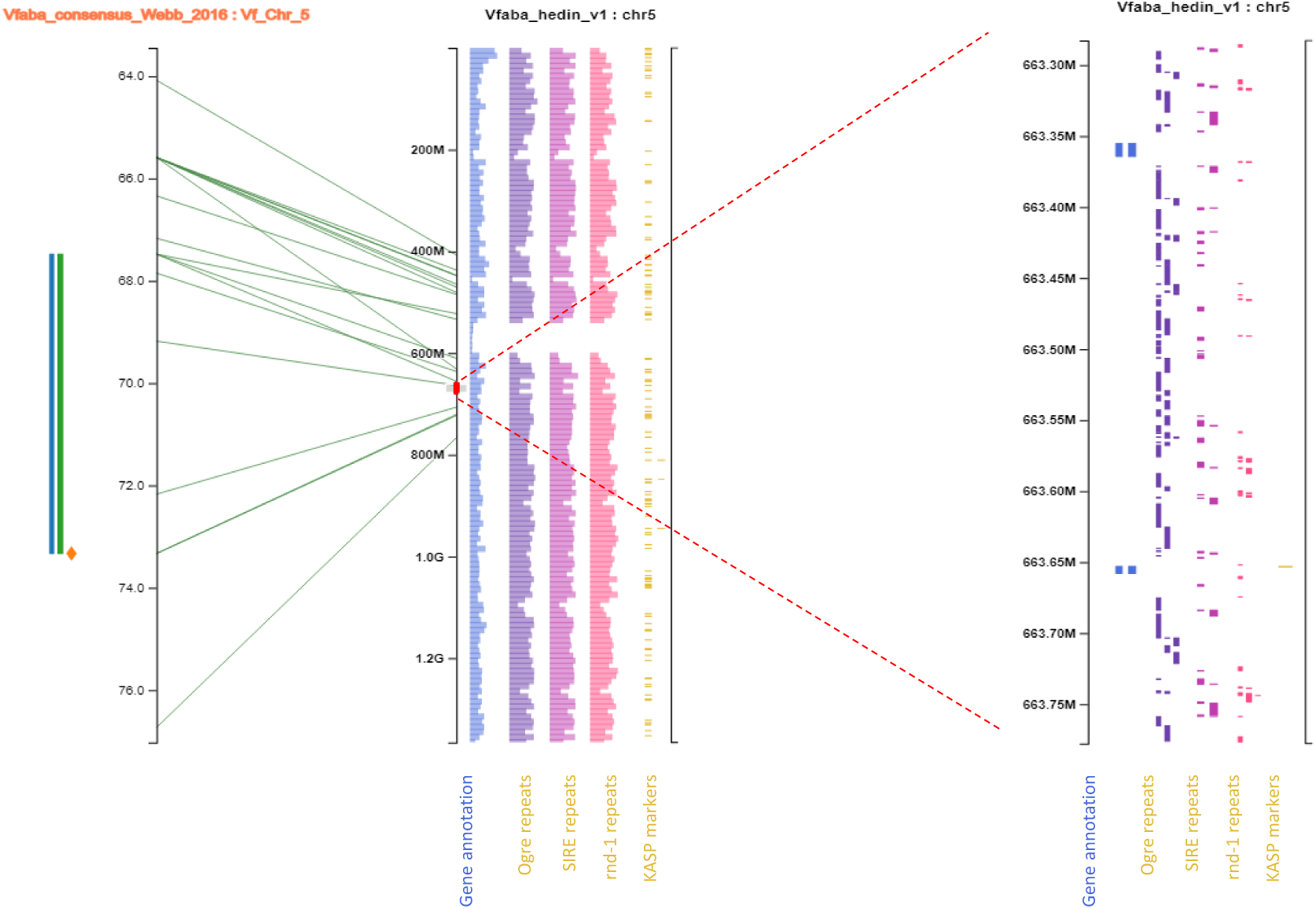
Pretzel (pulses.plantinformatics.io) visualisations integrating the consensus genetic map from Webb et al. (2016) together with 3 QTLs for frost tolerance from Sallam et al. (2016) (left axis), aligned to the Hedin/2 genome (middle and right axes). The gene annotation for Hedin/2 is shown together with the three most abundant repeat types (Ogre, SIRE and rnd-1). Pretzel enables rapid, interactive interrogation of multiple data types and integration of legacy knowledge with the new assemblies.

**Extended Data Figure 11.**
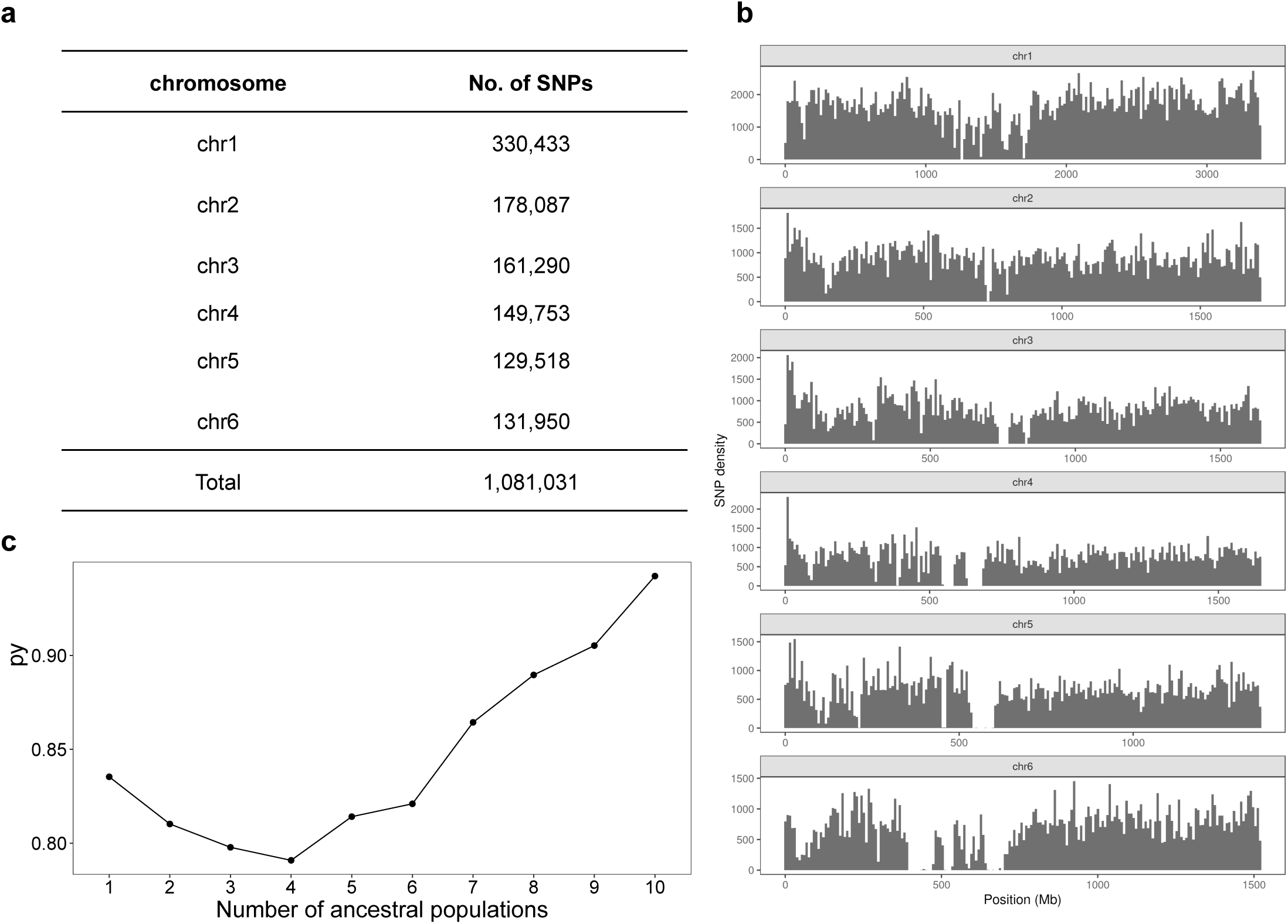
**a,** Summary of high-quality SNPs identified in each of the six chromosomes. **b,** Distribution of SNPs along each chromosome. **c,** The optimal K value estimated by Admixture analysis.

**Extended Data Figure 12.**
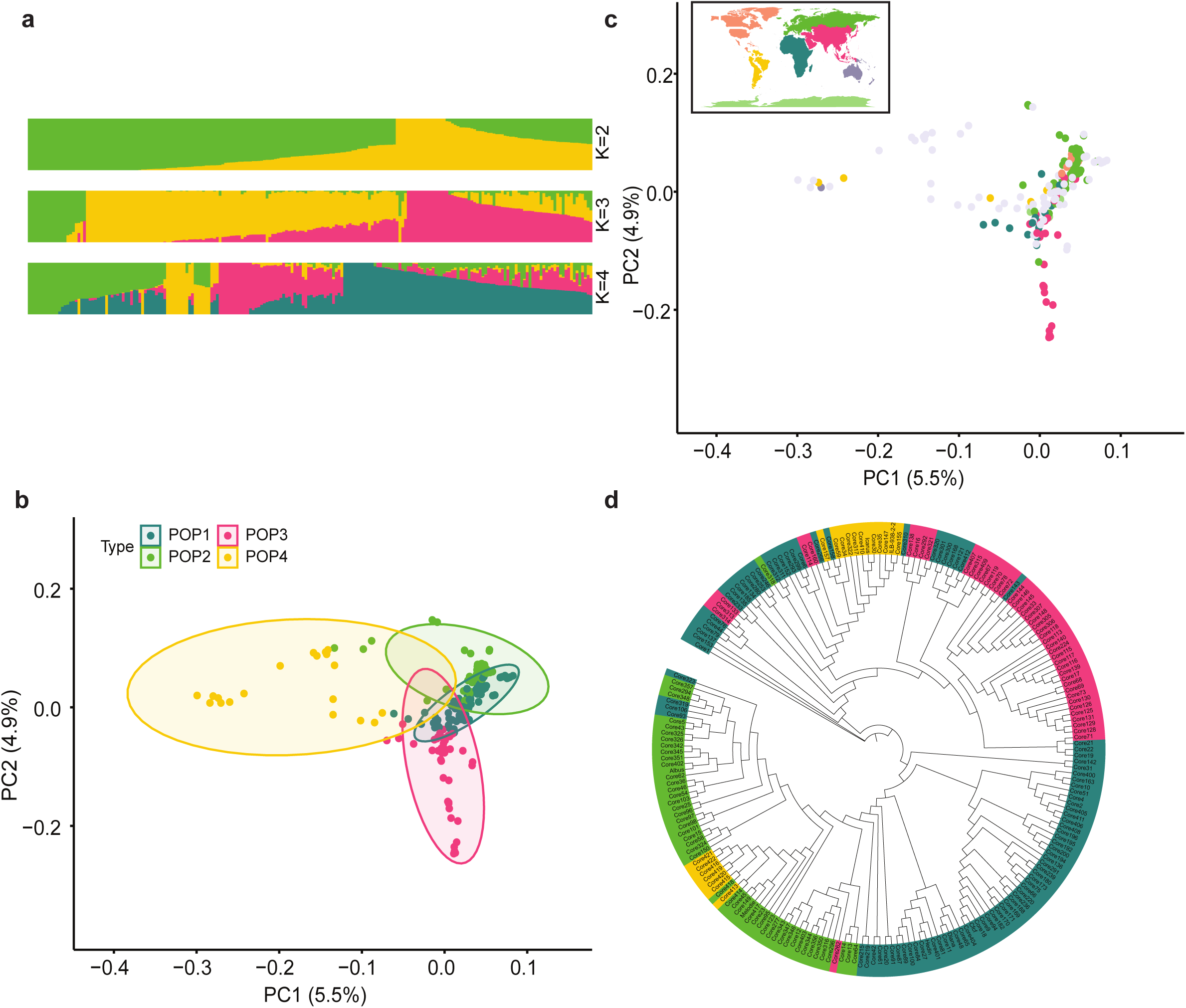
Genetic structure and association scan. **a,** Population structure grouped by ADMIXTURE **b,** Principal components analysis (PCA) of 197 accessions grouped with regard to subpopulation group. **c,** PCA of 197 accessions colored based on geographic origin. **d,** Neighbour joining (NJ) tree showing relationships among subpopulation group from 197 accessions.

**Extended Data Figure 13.**
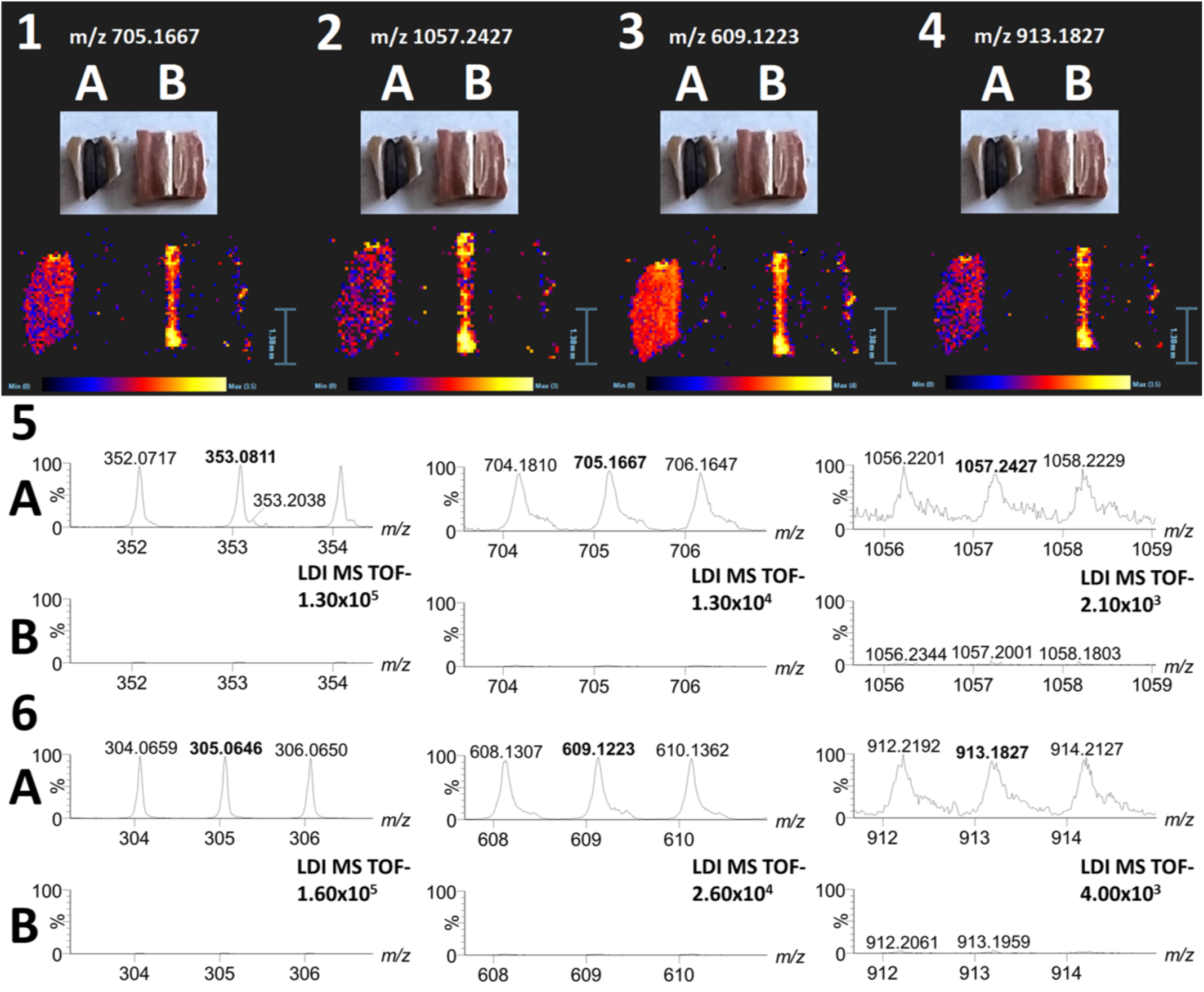
FIA-MS showing polymerization of phenolic compounds. Distribution of dimer and trimer of chlorogenic acid (1 and 2) and gallocatechin (3 and 4) on the surface of pigmented (A, Hedin/2) and non-pigmented (B, Tiffany) hila. Zoomed spectra of monomer, dimer and trimer of chlorogenic acid (5; signals at m/z 353.0811; 705.1667 and 1057.2427) and gallocatechin (6; signals at m/z 305.0646, 609.1223, 913.1827) collected from pigmented (A, Hedin/2) and non-pigmented (B, Tiffany) hila (the spectra showing particular signals are zoomed on the same intensity for both genotypes, A and B, e.g. the intensity 1.30×105 is set for spectra 5A and 5B, etc.), spectra are collected from the compact surface without the hilar groove area and edges of seed coat fragment.

**Extended Data Figure 14.**
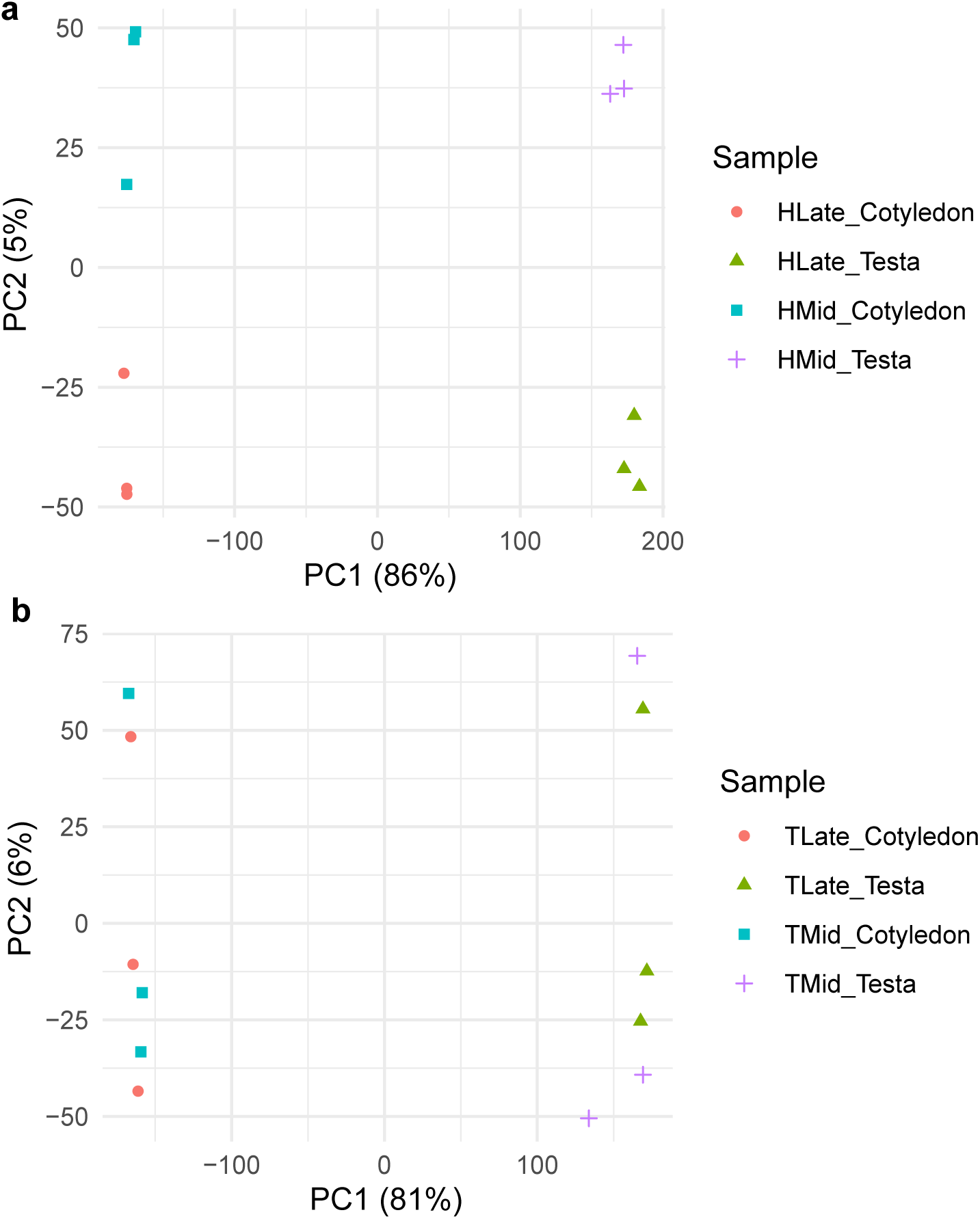
PCA of libraries showing tissue and developmental stage differentiation. a, Hedin/2 libraries. b, Tiffany libraries.

**Extended Data Figure 15.**
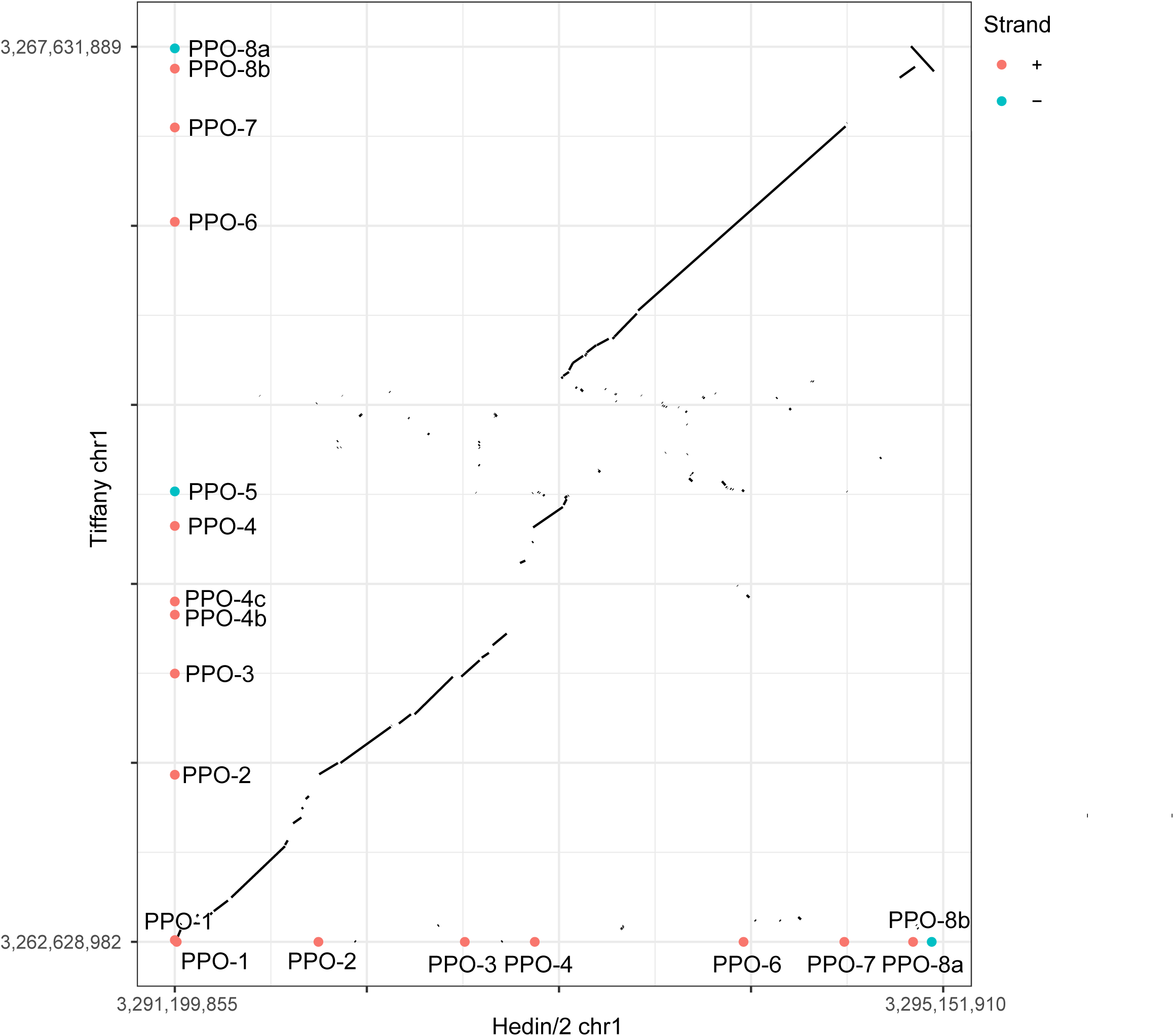
A dot plot alignment of PPO locus of Hedin/2 and Tiffany. The red and blue dots represent genes coloured according to strand.

**Extended Data Figure 16.**
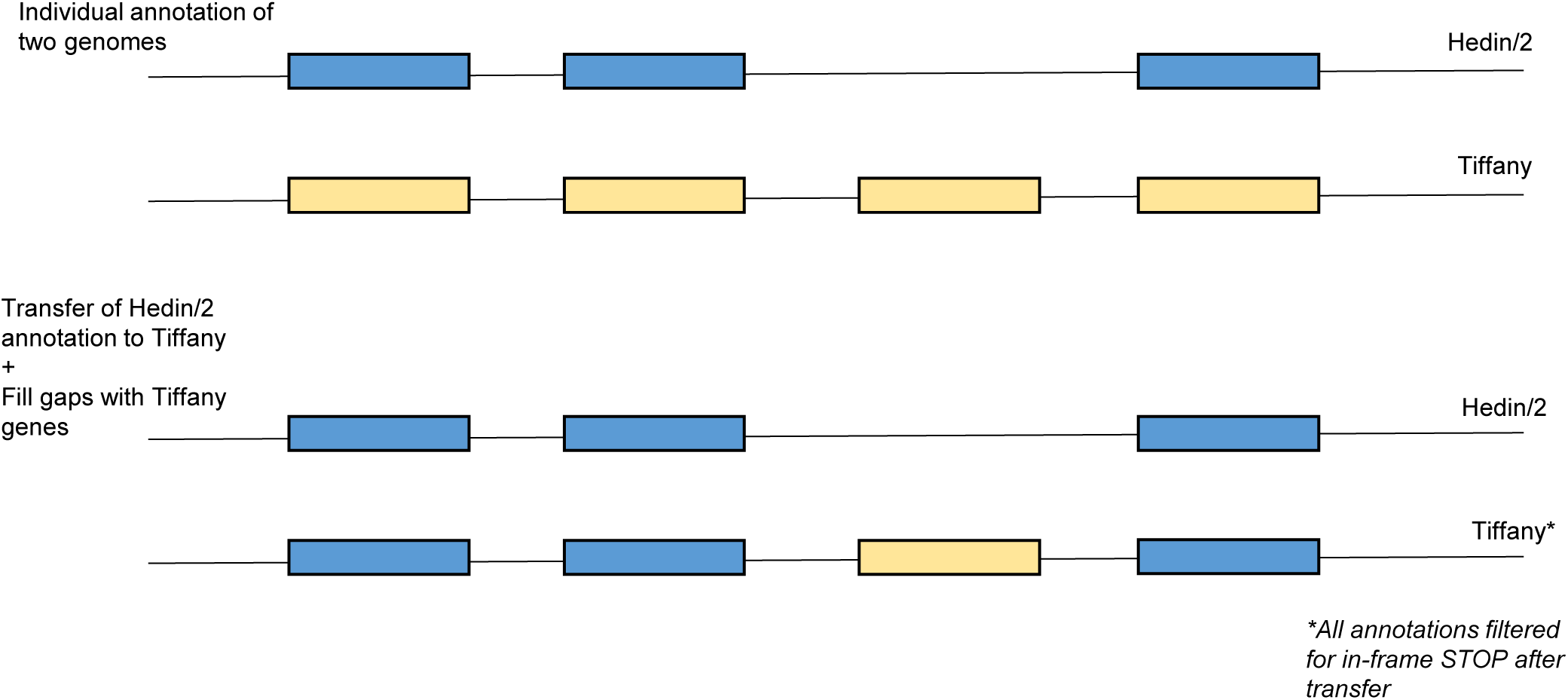
Principle behind individual annotations and ‘transfer and gap fill’ strategy.

**Extended Data Figure 17.**
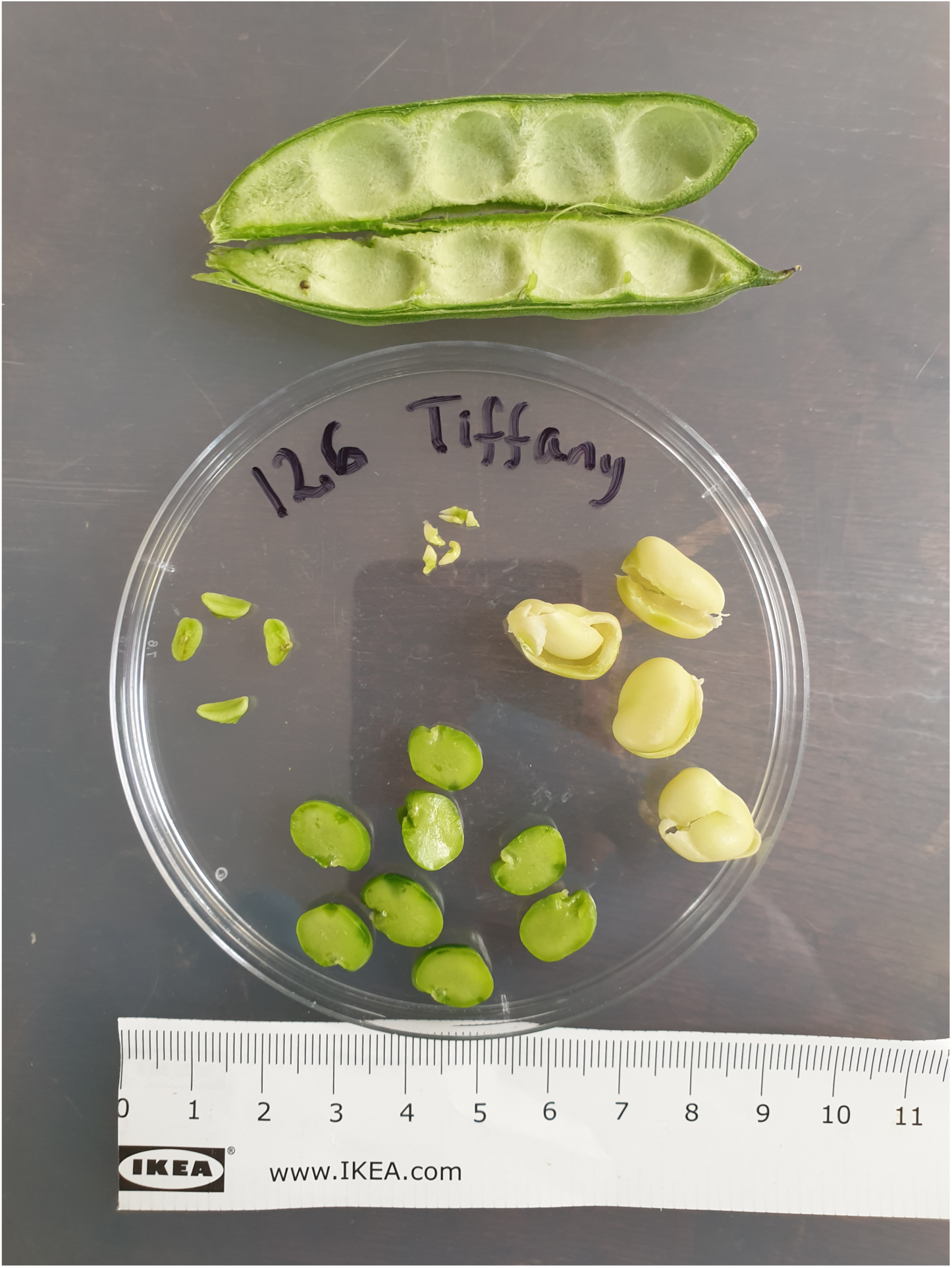
Dissected parts of a pod of cv ‘Tiffany’: pod wall, testa, cotyledon, funiculus and embryo axis.

**Extended Data Figure 18.**
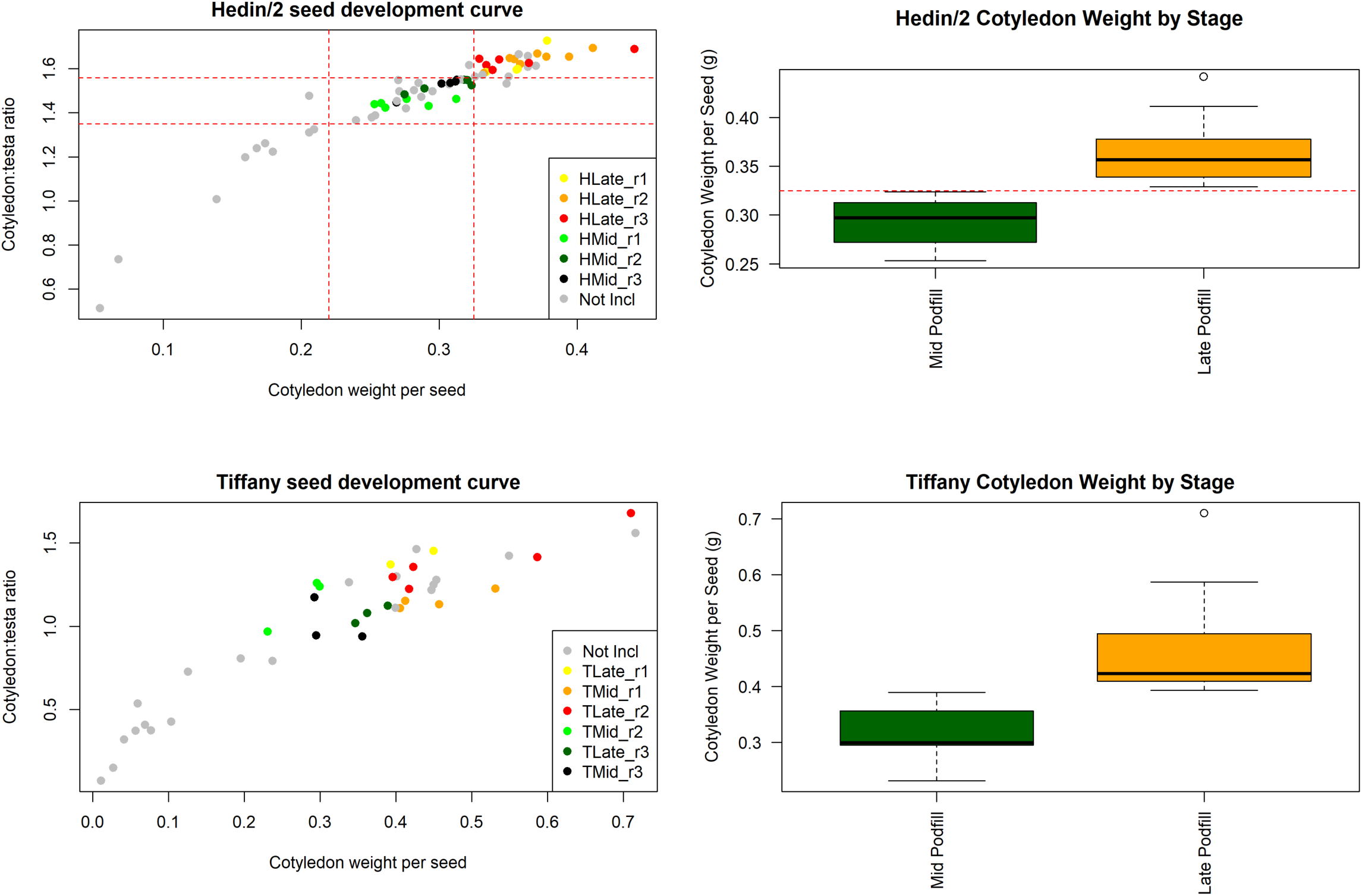
Composition and phenotypic differentiation of testa and cotyledon libraries from developmental pod-fill gradient.

## References

1. Hyland, J.J., Henchion, M., McCarthy, M. & McCarthy, S.N. The role of meat in strategies to achieve a sustainable diet lower in greenhouse gas emissions: A review. Meat Science 132, 189–195 (2017).

2. de Visser, C.L.M., Schreuder, R. & Stoddard, F. The EU’s dependency on soya bean import for the animal feed industry and potential for EU produced alternatives. OCL 21, D407 (2014).

3. Vranken, L., Avermaete, T., Petalios, D. & Mathijs, E. Curbing global meat consumption: Emerging evidence of a second nutrition transition. Environmental Science & Policy 39, 95–106 (2014).

4. Tanno, K.-i. & Willcox, G. The origins of cultivation of Cicer arietinum L. and Vicia faba L.: early finds from Tell el-Kerkh, north-west Syria, late 10th millennium b.p. Vegetation History and Archaeobotany 15, 197–204 (2006).

5. Caracuta, V. et al. The onset of faba bean farming in the Southern Levant. Scientific Reports 5, 14370 (2015).

6. Warsame, A.O., O’Sullivan, D.M. & Tosi, P. Seed Storage Proteins of Faba Bean (Vicia faba L): Current Status and Prospects for Genetic Improvement. Journal of Agricultural and Food Chemistry 66, 12617–12626 (2018).

7. Cernay, C., Pelzer, E. & Makowski, D. A global experimental dataset for assessing grain legume production. Scientific Data 3, 160084 (2016).

8. Herridge, D.F., Peoples, M.B. & Boddey, R.M. Global inputs of biological nitrogen fixation in agricultural systems. Plant and Soil 311, 1–18 (2008).

9. Bailes, E.J., Pattrick, J.G. & Glover, B.J. An analysis of the energetic reward offered by field bean (Vicia faba) flowers: Nectar, pollen, and operative force. Ecology and Evolution 8, 3161–3171 (2018).

10. Adhikari, K.N. et al. Conventional and Molecular Breeding Tools for Accelerating Genetic Gain in Faba Bean (Vicia Faba L.). Frontiers in Plant Science 12(2021).

11. Webb, A. et al. A SNP-based consensus genetic map for synteny-based trait targeting in faba bean (Vicia faba L.). Plant Biotechnology Journal 14, 177–185 (2016).

12. Björnsdotter, E. et al. VC1 catalyses a key step in the biosynthesis of vicine in faba bean. Nature Plants 7, 923–931 (2021).

13. Macas, J. & Neumann, P. Ogre elements — A distinct group of plant Ty3/gypsy-like retrotransposons. Gene 390, 108–116 (2007).

14. Macas, J. et al. In Depth Characterization of Repetitive DNA in 23 Plant Genomes Reveals Sources of Genome Size Variation in the Legume Tribe Fabeae. PLOS ONE 10, e0143424 (2015).

15. Chang, W., Jääskeläinen, M., Li, S.-p. & Schulman, A.H. BARE Retrotransposons Are Translated and Replicated via Distinct RNA Pools. PLOS ONE 8, e72270 (2013).

16. Fuchs, J., Strehl, S., Brandes, A., Schweizer, D. & Schubert, I. Molecular--cytogenetic characterization of the Vicia faba genome -- heterochromatin differentiation, replication patterns and sequence localization. Chromosome Research 6, 219–230 (1998).

17. Mascher, M. et al. A chromosome conformation capture ordered sequence of the barley genome. Nature 544, 427–433 (2017).

18. de Bruijn, F.J. The common symbiotic signaling pathway (CSSP or SYM). in The Model Legume Medicago truncatula 521–521 (2020).

19. Courty, P.E., Smith, P., Koegel, S., Redecker, D. & Wipf, D. Inorganic Nitrogen Uptake and Transport in Beneficial Plant Root-Microbe Interactions. Critical Reviews in Plant Sciences 34, 4–16 (2015).

20. Wipf, D., Krajinski, F., van Tuinen, D., Recorbet, G. & Courty, P.-E. Trading on the arbuscular mycorrhiza market: from arbuscules to common mycorrhizal networks. New Phytologist 223, 1127–1142 (2019).

21. De La Torre, A.R. et al. Insights into Conifer Giga-Genomes. Plant Physiology 166, 1724–1732 (2014).

22. Niu, S. et al. The Chinese pine genome and methylome unveil key features of conifer evolution. Cell 185, 204–217.e14 (2022).

23. Schmutz, J. et al. Genome sequence of the palaeopolyploid soybean. Nature 463, 178–183 (2010).

24. Cannon, S.B. et al. Multiple Polyploidy Events in the Early Radiation of Nodulating and Nonnodulating Legumes. Molecular Biology and Evolution 32, 193–210 (2014).

25. Ávila Robledillo, L. et al. Satellite DNA in Vicia faba is characterized by remarkable diversity in its sequence composition, association with centromeres, and replication timing. Scientific Reports 8, 5838 (2018).

26. Cossu, R.M. et al. LTR Retrotransposons Show Low Levels of Unequal Recombination and High Rates of Intraelement Gene Conversion in Large Plant Genomes. Genome Biology and Evolution 9, 3449–3462 (2017).

27. Bewick, A.J. & Schmitz, R.J. Gene body DNA methylation in plants. Current Opinion in Plant Biology 36, 103–110 (2017).

28. Brenet, F. et al. DNA Methylation of the First Exon Is Tightly Linked to Transcriptional Silencing. PLOS ONE 6, e14524 (2011).

29. Wang, L. et al. DNA methylome analysis provides evidence that the expansion of the tea genome is linked to TE bursts. Plant Biotechnology Journal 17, 826–835 (2019).

30. Barchi, L. et al. Single Primer Enrichment Technology (SPET) for High-Throughput Genotyping in Tomato and Eggplant Germplasm. Frontiers in Plant Science 10(2019).

31. Khazaei, H. et al. Flanking SNP markers for vicine–convicine concentration in faba bean (Vicia faba L.). Molecular Breeding 35, 38 (2015).

32. Balarynová, J. et al. The loss of polyphenol oxidase function is associated with hilum pigmentation and has been selected during pea domestication. New Phytologist 235, 1807–1821 (2022).

33. Gopher, A. et al. Breaking Ground: Plant Domestication in the Neolithic Levant : the "core-area One-event" Model, (Emery and Claire Yass Publications in Archaeology, The Institute of Archaeology, Tel Aviv University, 2021).

34. Scarborough, J. Beans, Pythagoras, Taboos, and Ancient Dietetics. The Classical World 75, 355–358 (1982).

35. Cubero, J.I. Taxonomy, Distribution and Evolution of the Faba Bean and its Mild Relatives. in Genetic Resources and Their Exploitation — Chickpeas, Faba beans and Lentils (eds. Witcombe, J.R. & Erskine, W.) 131–144 (Springer Netherlands, Dordrecht, 1984).

## Reference (online method)

1. Cheng, H., Concepcion, G.T., Feng, X., Zhang, H. & Li, H. Haplotype-resolved de novo assembly using phased assembly graphs with hifiasm. Nature methods 18, 170–175 (2021).

2. Li, H. Minimap2: pairwise alignment for nucleotide sequences. Bioinformatics 34, 3094–3100 (2018).

3. Carrillo-Perdomo, E. et al. Development of new genetic resources for faba bean (Vicia faba L.) breeding through the discovery of gene-based SNP markers and the construction of a high-density consensus map. Scientific reports 10, 1–14 (2020).

4. Monat, C. et al. TRITEX: chromosome-scale sequence assembly of Triticeae genomes with open-source tools. Genome biology 20, 1–18 (2019).

5. Ávila Robledillo, L. et al. Satellite DNA in Vicia faba is characterized by remarkable diversity in its sequence composition, association with centromeres, and replication timing. Scientific reports 8, 1–11 (2018).

6. Martin, M. Cutadapt removes adapter sequences from high-throughput sequencing reads. *EMBnet*. journal 17, 10–12 (2011).

7. Li, H. et al. The sequence alignment/map format and SAMtools. Bioinformatics 25, 2078–2079 (2009).

8. Doležel, J., Greilhuber, J. & Suda, J. Estimation of nuclear DNA content in plants using flow cytometry. Nature protocols 2, 2233–2244 (2007).

9. Doležel, J., Sgorbati, S. & Lucretti, S. Comparison of three DNA fluorochromes for flow cytometric estimation of nuclear DNA content in plants. Physiologia plantarum 85, 625–631 (1992).

10. Dolezel, J. Nuclear DNA content and genome size of trout and human. Cytometry Part A 51, 127–128 (2003).

11. Marçais, G. & Kingsford, C. A fast, lock-free approach for efficient parallel counting of occurrences of k-mers. Bioinformatics 27, 764–770 (2011).

12. Sun, H., Ding, J., Piednoël, M. & Schneeberger, K. findGSE: estimating genome size variation within human and Arabidopsis using k-mer frequencies. Bioinformatics 34, 550–557 (2018).

13. Sedlazeck, F.J. et al. Accurate detection of complex structural variations using single-molecule sequencing. Nature methods 15, 461–468 (2018).

14. Simão, F.A., Waterhouse, R.M., Ioannidis, P., Kriventseva, E.V. & Zdobnov, E.M. BUSCO: assessing genome assembly and annotation completeness with single-copy orthologs. Bioinformatics 31, 3210–3212 (2015).

15. Mapleson, D., Garcia Accinelli, G., Kettleborough, G., Wright, J. & Clavijo, B.J. KAT: a K-mer analysis toolkit to quality control NGS datasets and genome assemblies. Bioinformatics 33, 574–576 (2017).

16. Jiang, T. et al. Long-read-based human genomic structural variation detection with cuteSV. Genome biology 21, 1–24 (2020).

17. Roach, M.J., Schmidt, S.A. & Borneman, A.R. Purge Haplotigs: allelic contig reassignment for third-gen diploid genome assemblies. BMC bioinformatics 19, 1–10 (2018).

18. Alonge, M., et al. Automated assembly scaffolding elevates a new tomato system for high-throughput genome editing. BioRxiv (2021).

19. Lin, H.-N. & Hsu, W.-L. GSAlign: an efficient sequence alignment tool for intra-species genomes. BMC genomics 21, 1–10 (2020).

20. Flynn, J.M. et al. RepeatModeler2 for automated genomic discovery of transposable element families. Proceedings of the National Academy of Sciences 117, 9451–9457 (2020).

21. Brůna, T., Hoff, K.J., Lomsadze, A., Stanke, M. & Borodovsky, M. BRAKER2: automatic eukaryotic genome annotation with GeneMark-EP+ and AUGUSTUS supported by a protein database. NAR genomics and bioinformatics 3, lqaa108 (2021).

22. Dobin, A. et al. STAR: ultrafast universal RNA-seq aligner. Bioinformatics 29, 15–21 (2013).

23. Dobin, A. & Gingeras, T.R. Mapping RNA - seq reads with STAR. Current protocols in bioinformatics 51, 11.14. 1–11.14.19 (2015).

24. Kriventseva, E.V. et al. OrthoDB v10: sampling the diversity of animal, plant, fungal, protist, bacterial and viral genomes for evolutionary and functional annotations of orthologs. Nucleic acids research 47, D807–D811 (2019).

25. Björnsdotter, E. et al. VC1 catalyses a key step in the biosynthesis of vicine in faba bean. Nature plants 7, 923–931 (2021).

26. Wu, T.D. & Watanabe, C.K. GMAP: a genomic mapping and alignment program for mRNA and EST sequences. Bioinformatics 21, 1859–1875 (2005).

27. Liao, Y., Smyth, G.K. & Shi, W. featureCounts: an efficient general purpose program for assigning sequence reads to genomic features. Bioinformatics 30, 923–930 (2014).

28. Lyu, J.I. et al. Unraveling the complexity of faba bean (Vicia faba L.) transcriptome to reveal cold-stress-responsive genes using long-read isoform sequencing technology. Scientific reports 11, 1–13 (2021).

29. Quinlan, A.R. & Hall, I.M. BEDTools: a flexible suite of utilities for comparing genomic features. Bioinformatics 26, 841–842 (2010).

30. Li, P. et al. RGAugury: a pipeline for genome-wide prediction of resistance gene analogs (RGAs) in plants. BMC genomics 17, 1–10 (2016).

31. Kang, Y.-J. et al. CPC2: a fast and accurate coding potential calculator based on sequence intrinsic features. Nucleic acids research 45, W12–W16 (2017).

32. Shumate, A. & Salzberg, S.L. Liftoff: accurate mapping of gene annotations. Bioinformatics 37, 1639–1643 (2021).

33. Baggerly, K.A., Deng, L., Morris, J.S. & Aldaz, C.M. Differential expression in SAGE: accounting for normal between-library variation. Bioinformatics 19, 1477–1483 (2003).

34. Benjamini, Y. & Hochberg, Y. Controlling the false discovery rate: a practical and powerful approach to multiple testing. Journal of the Royal statistical society: series B (Methodological) 57, 289–300 (1995).

35. Altschul, S.F. et al. Gapped BLAST and PSI-BLAST: a new generation of protein database search programs. Nucleic acids research 25, 3389–3402 (1997).

36. Li, L., Stoeckert, C.J. & Roos, D.S. OrthoMCL: identification of ortholog groups for eukaryotic genomes. Genome research 13, 2178–2189 (2003).

37. Guindon, S., Delsuc, F., Dufayard, J.-F. & Gascuel, O. Estimating maximum likelihood phylogenies with PhyML. in Bioinformatics for DNA sequence analysis 113–137 (Springer, 2009).

38. Yang, Z. PAML: a program package for phylogenetic analysis by maximum likelihood. Computer applications in the biosciences 13, 555–556 (1997).

39. Wang, Y. et al. MCScanX: a toolkit for detection and evolutionary analysis of gene synteny and collinearity. Nucleic acids research 40, e49–e49 (2012).

40. Hasegawa, M., Kishino, H. & Yano, T.-a. Dating of the human-ape splitting by a molecular clock of mitochondrial DNA. Journal of molecular evolution 22, 160–174 (1985).

41. Zhang, Z. et al. KaKs_Calculator: calculating Ka and Ks through model selection and model averaging. Genomics, proteomics & bioinformatics 4, 259–263 (2006).

42. Ullrich, K.K. CRBHits: From Conditional Reciprocal Best Hits toCodon Alignments and Ka/Ks in R. The Journal of Open Source Software 5(2020).

43. Qiao, X. et al. Gene duplication and evolution in recurring polyploidization– diploidization cycles in plants. Genome biology 20, 1–23 (2019).

44. Haas, B.J., Delcher, A.L., Wortman, J.R. & Salzberg, S.L. DAGchainer: a tool for mining segmental genome duplications and synteny. Bioinformatics 20, 3643–3646 (2004).

45. Emms, D.M. & Kelly, S. OrthoFinder: phylogenetic orthology inference for comparative genomics. Genome biology 20, 1–14 (2019).

46. Balarynova, J. et al. The loss of polyphenol oxidase function is associated with hilum pigmentation and has been selected during pea domestication. New Phytol 235, 1807–1821 (2022).

47. Zablatzká, L., Balarynová, J., Klčová, B., Kopecký, P. & Smýkal, P. Anatomy and Histochemistry of Seed Coat Development of Wild (Pisum sativum subsp. elatius (M. Bieb.) Asch. et Graebn. and Domesticated Pea (Pisum sativum subsp. sativum L.). International Journal of Molecular Sciences 22, 4602 (2021).

48. Krejčí, P. et al. Combination of electronically driven micromanipulation with laser desorption ionization mass spectrometry – The unique tool for analysis of seed coat layers and revealing the mystery of seed dormancy. Talanta 242, 123303 (2022).

49. Warsame, A.O., Michael, N., O’Sullivan, D.M. & Tosi, P. Seed Development and Protein Accumulation Patterns in Faba Bean (Vicia faba, L.). Journal of Agricultural and Food Chemistry 70, 9295–9304 (2022).

